# Neural signatures of impaired semantic contextualization in Autism Spectrum Disorder

**DOI:** 10.64898/2026.03.16.712048

**Authors:** Melissa Franch, Kalman A. Katlowitz, Elizabeth A. Mickiewicz, James L. Belanger, Raissa K. Mathura, Hanlin Zhu, Xinyuan Yan, Taha Ismail, Ana G. Chavez, Assia Chericoni, Danika Paulo, Eleonora Bartoli, Tomasz M. Frączek, Nicole R. Provenza, Sameer A. Sheth, Benjamin Y. Hayden

## Abstract

Social and communicative deficits are defining characteristics of autism spectrum disorder (ASD). Some theories suggest that these challenges, among other autistic traits, may arise from differences in predictive coding, or how the brain uses context to predict and interpret incoming information. This idea has the potential to link symptoms of autism to specific neurocomputational processes, and is especially promising for communication, whose impairment is a hallmark of ASD. Here we leveraged the ability of large language models (LLMs) to quantify semantic contextualization to analyze a unique dataset of responses from hippocampal neurons obtained during language listening in three mild-to-severe autistic individuals with comorbid epilepsy. Key elements of semantic coding were preserved in all three individuals with ASD: single-neuron response dynamics, representation of word-word semantic relationships, and patterns of context-dependent shifts in meaning. However, relative to controls, ASD resulted in reduced neural signatures of contextualization: (1) neuronal responses were aligned with earlier, less contextual layers of GPT-2, (2) ASD patients had lower effective dimensionality of the neural subspace predicting semantics, (3) neural representations of word meaning were less influenced by preceding context, and (4) neural signatures of lexical surprisal were reduced. Together, these results support theories of autism that emphasize impairments in contextualization, and highlight the power of LLMs as a tool for quantifying the computational basis of neurodevelopmental disorders.

## INTRODUCTION

Autism spectrum disorder (ASD) is characterized by differences in social communication, language use, and repetitive behavior, yet the neural mechanisms that give rise to these features remain incompletely understood (APA, 2013; Wachtel et al., 2024; Lord et al., 2018; Vogindroukas et al., 2022). A growing body of theory proposes that many autistic traits can be explained by altered predictive processing - differences in how the brain uses prior knowledge to anticipate incoming sensory and cognitive information (Pellicano and Burr 2012; Van de Cruys et al., 2014; Noel and Angelaki 2023; Lawson et al., 2014; Friston 2010; Chrysaitis and Series, 2022). Evidence supporting this theory comes from EEG, fMRI, and MEG, but not at level of individual neurons, where predictions must ultimately be generated (Schneider and Blank, 2026; Pellicano and Burr, 2012; Noel et al., 2021; Lawson et al., 2017; Grisoni et al., 2019; Schneebeli et al., 2022; Alamia et al., 2026; Sapey-Triomphe et al., 2023; Brodski-Guerniero et al., 2018; Qela et al., 2025; Gonzalez-Gadea et al., 2015; van Laarhoven et al., 2020; Gomez et al., 2014). Additionally, the theory remains controversial, in large part due to limitations in our ability to rigorously test it (Chrysaitis and Series, 2022; Lawson et al., 2014; Van de Cruys et al., 2014; Brock 2012; Van Boxtel and Lu, 2013). Moreover, it remains unclear how to extend these ideas to communication, which is a defining feature of autism. Indeed, language provides a particularly powerful domain in which to examine these mechanisms: comprehension depends on *contextualization*, the continuous integration of prior linguistic and world knowledge to constrain interpretation of words and meanings. If predictive computations are atypical in autism, then the neural representations that support semantic processing may differ not only in strength but in their structure and context sensitivity.

Understanding these differences at the circuit level is therefore essential for linking computational theories of autism to the neural basis of language and meaning. Over the past decade, computational approaches have increasingly been applied to behavioral and neural data in the study of psychiatric disease, giving rise to the field of computational psychiatry (Montague et al., 2012; Wang & Krystal, 2014; Huys et al., 2016; Adams et al., 2016; Friston et al., 2014). By formalizing cognition in quantitative, model-based terms, this framework has enabled researchers to uncover subtle alterations in learning, inference, and decision processes that are often invisible to traditional summary statistics or categorical diagnostic comparisons (Adams et al., 2016). Such methods have been particularly powerful in revealing latent structure in high-dimensional datasets, allowing small but systematic differences in cognitive computation to be detected at both the behavioral and neural levels (Kriegeskorte and Douglas, 2018). We see a parallel opportunity in the study of language and semantic representation in the brain: large language models (LLMs) provide richly structured, high-dimensional embeddings that capture fine-grained relationships among words and concepts, quantifying the geometry of semantic space (Mikolov et al., 2013; Devlin et al., 2018; Radford et al., 2019). When paired with neural recordings, these models make it possible to ask whether the organization of semantic representations differs across populations, not only in overall magnitude or selectivity, but in the structure of the representational space itself (Franch et al., 2025; Katlowitz et al., 2025; Zhu et al., 2026). In this sense, LLMs extend the logic of computational psychiatry into the domain of neurolinguistics: rather than testing for gross differences in activation, they enable the detection of systematic shifts in how linguistic information is encoded, generalized, and abstracted.

The hippocampus is well positioned to mediate these processes because it sits at the intersection of memory, context, and semantic knowledge (Eichenbaum 2017; Shohamy and Turk-Browne, 2013; Duff and Brown-Schmidt, 2012). Beyond its canonical role in episodic memory, converging evidence indicates that hippocampal circuits contribute to the representation of concepts and relational structure, supporting the binding of items to context and the flexible generalization of meaning across experiences (Vigano and Piazza, 2021; Schapiro et al., 2017; Whittington et al., 2020; Constantinescu et al., 2016). Neurophysiological recordings in humans and animals have shown that hippocampal neurons can encode abstract features, conceptual relationships, and task-relevant structure, while neuropsychological and imaging studies implicate the medial temporal lobe in aspects of language comprehension that depend on contextual integration and associative inference (Franch et al., 2025; Katlowitz et al., 2025; Zhu et al., 2026; Piai et al., 2016; Duff and Brown-Schmidt, 2012; Courellis et al., 2024; Kolibius et al., 2023; Karkowski et al., 2025; Quiroga et al., 2005; Binder et al., 2009; Dijksterhuis et al., 2024). Importantly, several lines of evidence suggest that hippocampal structure and function are atypical in autism, including differences in volume and connectivity (Aylward et al., 1999; Reinhardt et al., 2023; Bhamidimarri et al., 2026; Hashimoto et al., 2021;), altered activation during memory and social tasks (Banker et al., 2021; Dickstein et al., 2013; Gomez et al., 2014), and disruptions in relational learning (Minor et al., 2023; Hashimoto et al., 2021). Together, these findings raise the possibility that circuit-level differences in hippocampal computation could contribute to the distinctive profile of contextual and semantic processing observed in autism, motivating direct investigation of how meaning is represented within hippocampal neurons.

## RESULTS

### Hippocampal encoding of semantic and phonemic features in autism

Fourteen adult patients (seven males and seven females) listened to 27 minutes of speech (**Methods, Figure 1A-B**). Of these patients, three were autistic, whose language abilities ranged from fluent to nonverbal (2 males and 1 female, see **Extended Data Fig. 1** for clinical and cognitive assessments). The monologues consisted of a total of 4,128 words, of which 792 were unique. We collected responses of isolated single neurons in the hippocampus (HPC, n=385 neurons in the control cohort; n=91 in the ASD cohort, **Extended Data Fig. 2**). For this study all analyses are performed at the individual patient level. Many neurons in both patient groups showed a word-aligned ramping (**Figure 1C**). We defined word-evoked firing in a window lasting the duration of the word starting 80 ms after the onset of each word (to account for the approximate response latency; see **Methods** and Franch et al., 2025).

**Figure 1.**
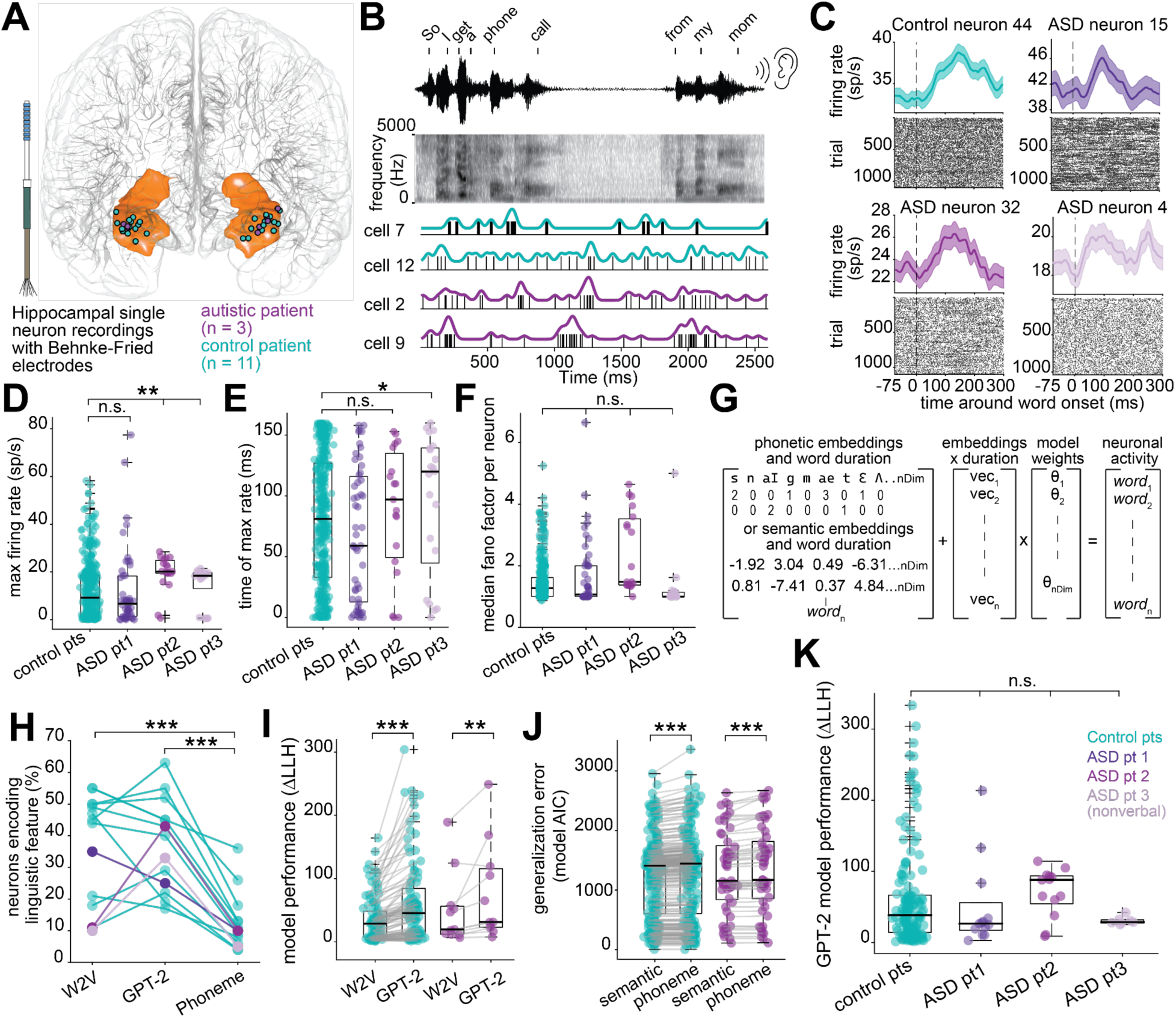
Preserved single neuron semantic and phonetic encoding in autism. **A,** Electrode recording location of hippocampal neurons from 11 control patients and 3 patients with autism. **B,** Continuous spiking activity from four example hippocampal neurons from control patient 7 and autistic patient 2 during words heard during the experiment, as shown in the spectrogram. **C,** Peri-stimulus time histograms and raster plots demonstrating neural responses to word onsets from four distinct hippocampal neurons in four different patients. Dashed lines represent word onset. Plots display mean firing rate ±SEM shaded. **D,** For each neuron (circle), the peak word-averaged firing rate following word onset. ASD2-3 had higher firing rates than controls (P < 0.01, two-sided randomization test on medians using 5,000 neuron-matched resamples from controls). **E,** For each neuron, the time after word onset that peak word-averaged firing rate occurred. ASD3 had a modest increase in latency relative to controls (P = 0.04, two-sided randomization test on medians using 5,000 neuron-matched resamples from controls). **F,** For each neuron, the median fano factor across all words that occurred at least ten times (n = 66 words). **G,** Regression equation used to fit an individual neuron’s spike count from word embeddings or phonetic embeddings, word duration, and their pairwise interactions. **H,** For each patient (circle), their percentage of neurons fit by the regression model in G using semantic embeddings from Word2Vec or GPT-2 layer 25 or IPA phonemic embeddings. The percentage of neurons fit by phonetic embeddings was significantly lower relative to semantic embedding models (P < 0.001, one-sided Wilcoxon signed-rank test). **I,** Log-likelihood improvements (ΔLLH; model performance) for fitted neurons (n = 109) from GPT-2 and Word2Vec semantic encoding models. Word2Vec performance is significantly lower than GPT-2 in control and autistic patients (P < 0.001, one-sided Wilcoxon signed-rank test). **J,** Model error (penalized Poisson AIC, EDF-based) per neuron for phonetic and semantic embedding models. AIC is consistently lower for semantic models (GPT-2 shown) across control and autistic patients (P < 0.001, Wilcoxon signed-rank test). **K,** Log-likelihood improvements (ΔLLH; model performance) for fitted neurons (n = 188 across all patients) from Poisson ridge regression predicting neuronal spike counts from the first 100 PCs of semantic (GPT-2 layer 25) embeddings using the equation in G. Autistic patient performance is comparable to controls (P = 0.15, 0.96. and 0.26 for ASD1-3 respectively, one-sided Monte Carlo subsampling test on medians using 5,000 neuron-matched resamples from controls).

Evoked firing rates in ASD1 were comparable to controls while firing rates in ASD2 and ASD3 were enhanced relative to our control cohort (p=0.31, 0.0005, and 0.004, respectively; Monte Carlo subsampling test; **Figure 1D**). Latency to peak was greater in ASD3 relative to controls, although only modestly so (increase in 40 ms to median latency, p = 0.041, **Figure 1E**). Fano factor was not statistically different in any of the patients (all p>0.05, **Figure 1F**).

To understand the neural representation of semantics and phonemics, we took an encoding model approach (Huth and Gallant, 2016; Goldstein et al., 2024; Goldstein, Wang, et al., 2023; Tang et al., 2023; Zada et al., 2024) that we and others have adapted for use in single neurons (Franch et al., 2025; Jamali et al., 2024;. Khanna et al., 2024). **Figure 1G**). Our regression approach controls for phonemic embeddings, word duration, and their interactions (cf. Khanna et al., 2024). We find robust semantic encoding in all patients using both Word2Vec and GPT-2 embeddings (**Figure 1H-J**). In both groups, the contextual (GPT-2) embedder fit more neurons than the non-contextual one (Word2Vec), and both semantic models fit many more neurons than the phonemic model (all p<0.001, **Figure 1H**). GPT-2 had significantly better fit than Word2Vec in both control and autistic patients (p<0.001 and p<0.01, one-sided Wilcoxon signed-rank test between ΔLLH of fitted neurons in each semantic model, **Figure 1I**). Additional controls also show that semantics remains the dominant driver of neural responses even when accounting for word frequency, word order in sentence, word order in clause, and number of opening and closing nodes (**Extended Data Figs. 3-4**). Semantic encoding model performance (ΔLLH) and explained variance (pseudo R²) did not differ between autistic and control patients, suggesting intact neural encoding of semantics in autism (median pseudo R² = 0.16, 0.14, 0.24, and 0.15 for control patients and ASD1-3 respectively; p = 0.15, 0.96. and 0.26 for ASD1-3 respectively, Monte Carlo subsampling test; **Figure 1K**). These findings indicate that single neuron encoding of basic linguistic features in autism is similar to that in controls.

### Altered semantic contextualization in ASD

Individual word meanings can be thought of as vectors in a high-dimensional semantic embedding space, where vector angle corresponds to semantic similarity (Mikolov et al., 2013; Devlin et al., 2018; Radford et al., 2019; **Figure 2A**). For example, the vector angle between ‘cat’ and ‘spoon’ will be larger than that for ‘cat’ and ‘tiger.’ When words appear in context, their meanings change - sometimes modestly, sometimes profoundly - a process known as contextualization. Contextual changes in word meaning are captured in LLMs such as GPT-2 (Radford et al., 2019).

**Figure 2.**
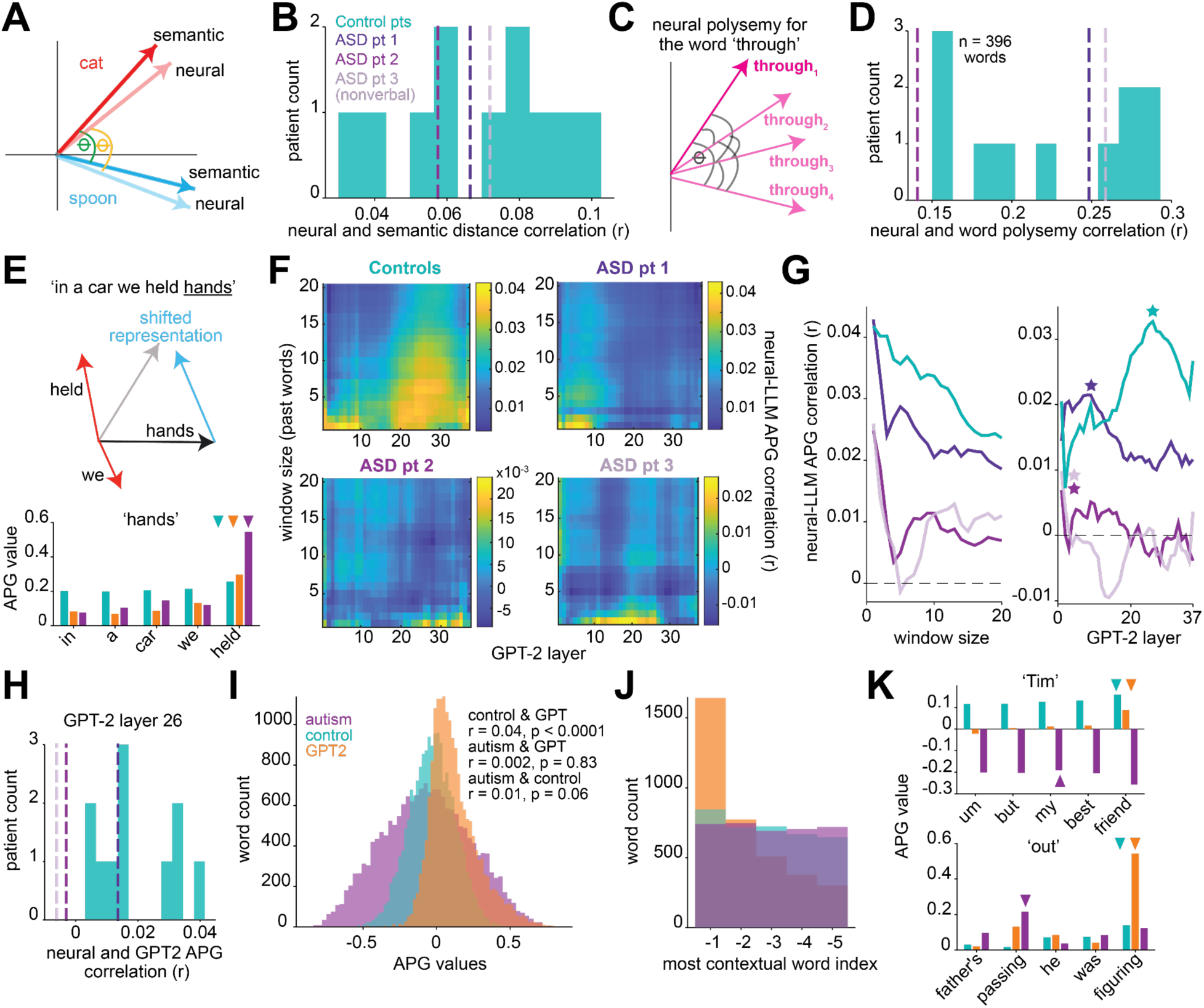
Reduced prior-word context integration in semantic neural coding in autism. **A,** We define semantic distance as the angular distance in a high dimensional space between two semantic vectors (embedding values for each word; green theta). We define neural distance in the analogous manner, where each word’s neural embedding is the vector of firing rates from all neurons for that word (yellow theta). **B,** For each patient, the Pearson correlation between neural and semantic embedding (GPT-2 layer 25) distances between all 8,514,001 word pairs (all p < 0.001). ASD correlations are shown as dashed lines to visualize their relation to controls. **C,** Schematic showing how we define polysemy as the median cosine distance in neural population or embedding space between different occurrences of the same word. **D,** For each patient, the Pearson correlation between neural and word polysemy (n = 396 words; all p < 0.001). ASD correlations are shown as dashed lines to visualize their relation to controls. **E,** Top - Vectorial diagram shows how adjacent words can modify the neural vectorial representations of words (e.g., “held” appearing in the context of “hands” shifts the firing rate vector for “hands” to have a more similar cosine distance to the vector for “held”; “hands” vector shifts from black to gray). Bottom - A real example sentence of the APG values computed for six words occurring before the word “hand” from GPT-2 or neural embeddings from each patient group (median across patients). Triangles indicate the most contextualizing word for “hands”. **F,** The correlation between neural and GPT-2 APG values computed for each GPT-2 layer (x-axis) and up to 20 previous words (y-axis). The top left plot shows the median correlation across all control patients. **G,** From the data in F, the neural-APG correlation values plotted across different window sizes (left) and GPT-2 layers (right). The teal line reflects median correlation across controls. Stars reflect the layer of maximum correlation; for visibility, stars for ASD2-3 (layer 1) are plotted slightly to the right. **H,** For each patient, the Pearson correlations between neural and GPT-2 APG values (n = 18,015 APG values) computed from layer 26 and for the previous five words. Autistic patient correlations are shown as dashed lines to visualize their relation to control patients. **I,** Histograms of 18,015 APG values computed separately from GPT-2 layer 26, control patient median, and ASD patient median. **J,** Histograms indicate which preceding word (1–5 words back) exerted the strongest contextual influence on the current word based on maximal APG values (n = 4,128 words). **K,** Real examples of APG values computed for the five words before “Tim” and “out” from GPT-2 layer 26 or neural embeddings from each patient group. Triangles indicate the most contextualizing word.

We found that, in both control and autistic patients, neural distance recapitulates semantic distance (all correlations p<0.001; **Figure 2B**). Indeed, autistic patients’ neural-semantic distance correlations did not differ from controls (p = 0.91,0.63, and 0.45, Crawford-Howell significance test; **Figure 2B**). This lack of a difference argues against the possibility that autistic patients have increased neural variability, such as may be caused by failure to attend to speech or to noisy internal representations.

We next considered how well control and autistic patient’s neural population codes were tracking the meaning of a word across contexts. For each word that occurred at least twice (n = 396 words), we quantified *polysemy* as the cosine distance between GPT-2 layer 25 embedding vectors across repeated occurrences (see **Methods;** Cruse etal., 1986; Landauer, 2001; Ethayarajh, 2019; **Figure 2C**). We computed a word’s *neural polysemy* as the median cosine distance between neural population responses to repeated occurrences of the same word. We then computed the correlation between neural polysemy and embedding-based polysemy across words (**Figure 2D**). In all patients, we find a positive correlation between neural polysemy and word polysemy (all p < 0.005; **Figure 2D**); autistic patients’ correlations were comparable to controls (p = 0.67, 0.18, and 0.54, Crawford-Howell significance test). These results demonstrate that core principles of semantic coding, semantic relationships and context-dependent shifts in meaning, are preserved in our autistic patients.

The specific strength of mappings between the past and current words in transformer-based LLMs is given by the ***attention pattern grid*** (**APG**, Vaswani et al., 2017; Vig, 2019). The APG is a data-derived vectorial representation of how strongly each prior word influences a given word, so the match between a neural and LLM-derived APG indexes how closely the brain implements contextualization (Katlowitz et al., 2025; **Figure 2E**). Here, we used this approach to determine whether semantic contextualization is altered in ASD.

We created 37 latent LLM APGs, one for the initial embedding and then each of the 36 layers of GPT-2 and compared each one to our neural APG from control and autistic patients (see **Methods**; **Figure 2F**). We tested correlations across a range of contextual windows (1-20 words) for each layer. In all three autistic patients, we find reduced contextualization at all window sizes except the shortest ones and at all layers. In other words, in the autistic patients, lexical contextualization is preserved, but with reduced magnitude at more than a few words in the past.

These effects can be observed even more clearly when considering the graph marginals (**Figure 2G**). Autistic patients showed a sharp drop in correlation immediately after the preceding word, suggesting a more restricted temporal integration of contextual information (**Figure 2G-left**). In control patients, neural-APG correlation peaked at layer 26, a late transformer layer associated with deep contextual semantic integration, whereas autistic patients showed peak attention alignment at earlier, less contextualized layers (stars - layers 8, 1, and 1 for ASD1-3 respectively, **Figure 2G-right;** Ethayarajh 2019; He et al., 2024).

ASD2 and ASD3 had a non-significant negative correlation with APG values from GPT-2 layer 26, while ASD1 and control patients had significant positive APG correlations with this layer (p<0.05, Pearson correlation; **Figure 2G-right and Figure 2H**). We pooled median APG values from up to five preceding words across all words within each group, generating one distribution for controls and one for autistic patients. (**Figure 2I**). We find that ASD patients have a broad distribution compared to both the control-neural and LLM APG values (**Figure 2I**). Indeed, the control patient APG values are positively correlated with LLM APG but the autistic patient’s APGs are not significantly correlated with either the control patients or LLM (**Figure 2I**; results are comparable even if only autistic patients 2-3 are used; **Extended Data Fig. 5**). We also find that both the LLM and control patients most frequently assign the *highest* contextual weight to the immediately preceding word (i.e., ‘*red*’ as the most contextualizing word in ‘*red balloon*’), whereas autistic patients show a broader distribution, with the most contextualizing weights often extending to more distant words up to five words back (**Figure 2J-K**). Together, these results indicate that neural contextual weighting in autistic patients is less precisely aligned with the statistical structure captured by the LLM and observed in control patients. Even when the same prior word exerts the strongest contextual influence, neural activity in autism shows differences in the sign and/or magnitude of the corresponding APG value relative to both LLMs and controls, reflecting altered scaling of contextual semantic influence rather than a complete loss of sensitivity (**Figure 2E-bottom and Figure 2K**).

### Reduced dimensionality of the predictive neural subspace for semantic coding in autism

We next asked how the hierarchical structure of semantic representations across GPT-2 layers is reflected in neural coding and whether it differs in autism. First, we calculated semantic embeddings from each GPT-2 layer and regressed them to neural activity as shown in Figure 1G. We find that ASD1 and ASD3 show a lower proportion of neurons fit by the model across GPT-2 layers, although overall model performance remains comparable to controls (**Figure 3A**). Importantly, single neuron responses in each autistic patient aligned *best* with semantic embeddings from earlier, less contextual layers of GPT-2 (layers 9, 15, and 12), whereas neurons from control patients aligned best with layer 27 (**Figure 3B**; range: 15-37). Embeddings from earlier layers are more strongly driven by lexical identity, word frequency, and part of speech, whereas layers 24 and beyond increasingly reflect semantic similarity and predictive probability (Ethayarajh 2019; He et al., 2024). To quantify significance, we split the data into ten contiguous segments and ran cross-validated regressions. Across models, autistic patients showed a consistent bias toward earlier GPT-2 layers relative to controls (p < 0.001, Wilcoxon rank-sum; **Figure 3B**). Notably, the association between late layers and semantic encoding is well-established in allistic adults, so the autistic data represent an exception to a larger pattern (Caucheteux etal., 2022; Goldstein, Ham, et al., 2022; Schrimpf et al., 2021).

**Figure 3.**
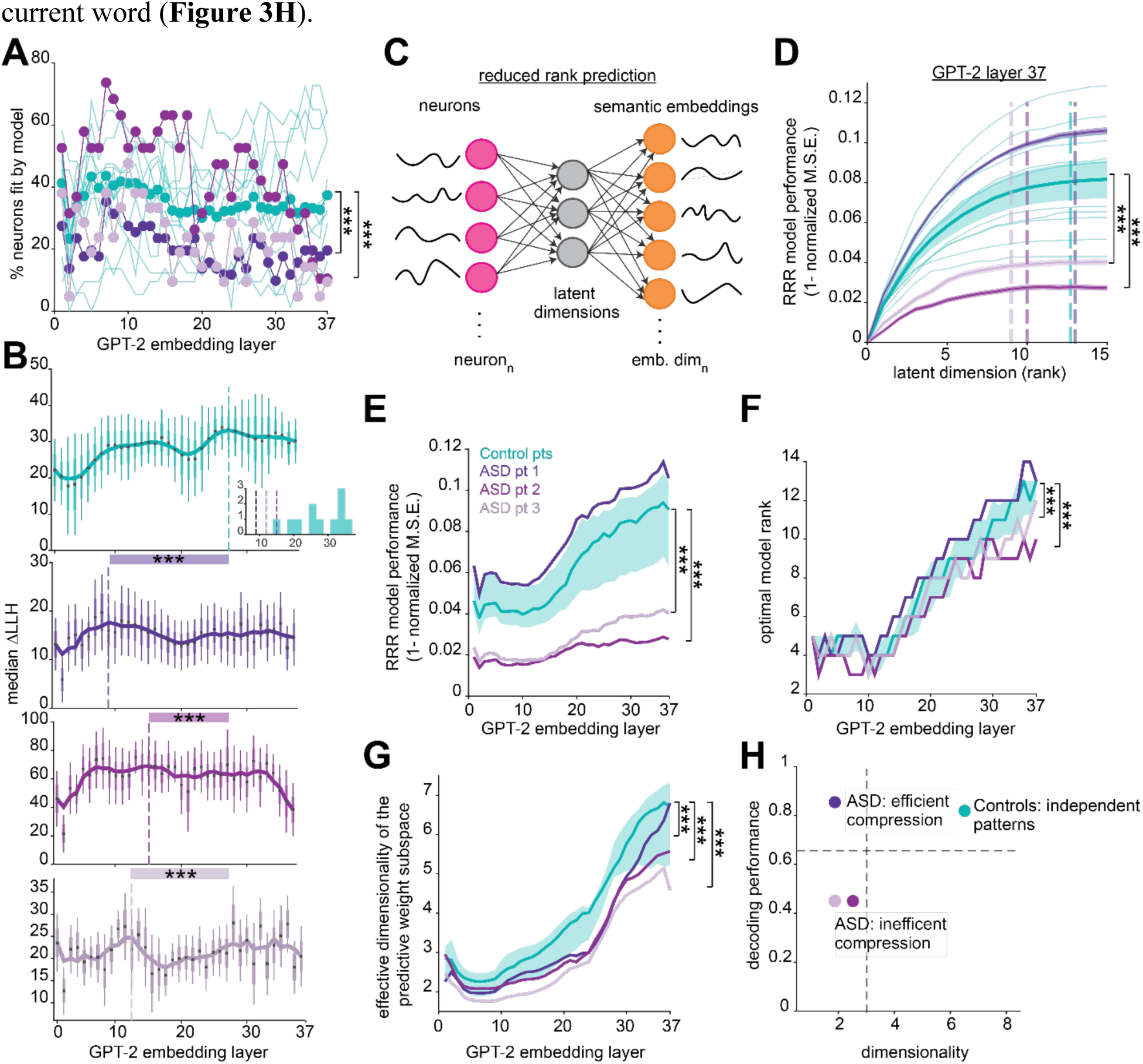
Reduced dimensionality of semantic coding in autism. **A,** Percentage of neurons fit by linear regression models using the first 100 PCs of semantic embeddings from each layer of GPT-2. Light teal lines show individual control patients; dark teal circles denote the median across controls. Two autistic patients have fewer neurons fit by the model relative to controls (P< 0.001, one-sided Wilcoxon signed-rank test comparing patient percentages to control median). **B,** The median model performance (ΔLLH) across neurons for each patient and layer (median and SEM). The bolded line represents the median smoothed with a gaussian of width 5 and dashed lines indicate the maximum of the fitted line. The insert shows the GPT-2 layer with best performance for each patient. Each autistic patient has peak performance at earlier layers (layers 9, 15, and 12) of the model compared to controls (layer 27; P < 0.001, Wilcoxon rank-sum test comparing each autistic patient’s values to controls across models). **C,** Schematic of reduced-rank regression (RRR) predicting semantic embeddings from neural population activity by projecting activity onto a low-dimensional set of predictive latent dimensions that explain variance in the predicted embedding space. Schematic adapted from those in Wu and Pillow, 2025. **D,** Model performance across 15 latent dimensions (ranks) from RRR predicting 30-dimensional (top 30 PCs) GPT-2 layer 37 semantic embeddings from neural firing rate responses to words. Models were fit separately for each patient, then averaged across control patients. Lines represent mean performance ±SEM. Model performance was significantly lower for two autistic patients compared to controls (P < 0.001, one-sided permutation test). Dashed lines indicate the number of dimensions for optimal performance; rank = 13 for controls and ASD1, rank = 10 and 9 for ASD2-3 respectively. **E,** For each GPT-2 layer, optimal RRR model performance is plotted. The dark teal line shows the median across control patients, and shading reflects the interquartile range. Model performance was significantly lower for ASD2-3 compared to controls (P < 0.001, one-sided permutation test). **F,** For each GPT-2 layer, the rank of optimal model performance is plotted. The dark teal line shows the median across control patients, and shading reflects the interquartile range. Model rank for ASD2-3 was consistently lower relative to controls across layers (P < 0.001, one-sided Wilcoxon signed-rank test). **G,** For each GPT-2 layer, the dimensionality of neuron-weight patterns used for semantic prediction is plotted. The dark teal line shows the control median, and shading indicates the interquartile range. Effective dimensionality of each autistic patient was significantly lower than control patients across layers (P < 0.001, one-sided permutation test). **H,** Predictive neural dimensionality reflects a trade-off between compression and representational coverage. A lower-dimensional predictive subspace can support strong performance when task-relevant signals align with dominant latent axes, whereas a higher-dimensional predictive subspace distributes variance across multiple independent axes, enabling greater flexibility and generalization.

We next investigated the geometry of semantic coding in autism. Task-related population activity often lies in a low-dimensional subspace relative to the number of recorded neurons or task variables (Gao et al., 2017; Cunningham and Yu, 2014; Ebitz and Hayden, 2021). To identify the low-dimensional neural subspace that predicts semantic structure, we applied cross-validated reduced rank regression (RRR) to map neural activity onto GPT-2 semantic embeddings (**Figure 3C**; using 30-dimensional embeddings; results were consistent across models using different numbers of PCs; **Methods**; Izenman 1975; Franch et al., 2025).

We quantified model performance as the cross-validated variance explained (1 – NSE) across predictive subspace dimensionalities (ranks = 0 - 30). All models showed near-zero performance at rank 0 (predicting the mean), confirming an unbiased baseline. Across ranks, predictive performance increased monotonically before reaching a plateau. ASD2-3 showed significantly lower model performance across ranks compared to each control patient, whereas ASD1 performed comparably to controls (p < 0.001, one-sided permutation test; **Figure 3D**). This pattern was consistent across all GPT-2 layers (**Figure 3E**). Across patients, decoding performance was highest for embeddings from mid-to-late GPT-2 layers; however, performance in ASD2-3 remained significantly lower than in controls and ASD1 (p<0.001, one-sided permutation test; **Figure 3E**). In other words, neural decoding of semantics was reduced in two autistic patients compared to controls.

We next examined the latent structure of the predictive subspace in control and autistic patients. Model rank in RRR corresponds to the number of orthogonal latent neural dimensions contributing to the predictive mapping from neural activity to the embedding space. Thus, rank provides a measure of how many independent population axes are required to support semantic prediction. Optimal model rank was defined as the smallest rank achieving performance within one standard error of peak cross-validated accuracy. For Layer 37, ASD2-3 exhibited lower optimal ranks than controls and ASD1 (dashed lines, **Figure 3D**). Across individuals, optimal rank increased for higher GPT-2 layers (from ∼layer 11 onward), indicating that deeper, more contextual language representations required a more distributed neural predictive subspace (**Figure 3F**). However, optimal rank remained significantly reduced across GPT-2 layers in ASD2-3, suggesting that semantic prediction in these individuals relied on fewer independent neural population dimensions (p<0.001, one-sided Wilcoxon signed-rank test; **Figure 3F**).

To determine how predictive variance is distributed across dimensions, we quantified the effective dimensionality of the predictive weight subspace using the participation ratio (**Methods**). This measures how evenly predictive variance is spread across latent axes. Intuitively, effective dimensionality distinguishes between a subspace in which predictive variance is concentrated along a small number of dominant dimensions and one in which variance is more uniformly distributed across many axes. For each patient, effective dimensionality increased across GPT-2 layers, indicating that predictive variance became more evenly distributed for higher-level semantic embeddings (**Figure 3G**). Notably, effective dimensionality was significantly reduced in all autistic patients across layers (p < 0.001, one-sided permutation test; **Figure 3G**), indicating that semantic prediction relied on a more compressed predictive geometry, with variance concentrated along a limited set of dominant neural axes.

These findings raise the possibility that autism may be associated with a bias toward lower-dimensional predictive representations of semantic information (**Figure 3H**). This compression is not inherently detrimental, as demonstrated by ASD1, whose decoding performance matched that of controls, suggesting that task-relevant semantic information can be preserved within a lower-dimensional subspace. However, when critical semantic variance is not adequately captured within this compressed predictive geometry, decoding performance may be impaired, as observed in ASD2-3. In contrast, control patients showed higher-dimensional predictive subspaces, which may allow the brain to simultaneously represent multiple aspects of a word’s meaning - such as its predictability, relationship to surrounding words, and role within the broader discourse - resulting in a richer and more detailed semantic representation of the current word (**Figure 3H**).

### Altered population-level coding of word surprisal in autism

Predictive coding theories propose that neural activity reflects the discrepancy between expected and observed input (Rao and Ballard, 1999; Feldman and Friston, 2010). In language, prediction error can be quantified using word surprisal, which measures how unexpected a word is given its context (Levy, 2008; Weissbart et al., 2020). We computed word-by-word surprisal values (see **Methods**). To isolate neural effects of surprisal, we residualized firing rates with respect to log word frequency and word duration, as both variables correlate with surprisal (Pearson r = −0.49 and 0.42, respectively).

To assess how neural activity relates to word surprisal, we quantified the relationship between residual firing rates and surprisal for each neuron. Across neurons, this relationship was predominantly monotonic, with lower-than-expected firing rates for less surprising words and higher residual firing rates for more surprising words (**Figure 4A**). On average, 54% of neurons in control patients and 62% in autistic patients were significantly correlated with word surprisal (**Figure 4B**; Spearman; FDR-corrected P < 0.05). Among these neurons, 73% and 78% in control and ASD patients, respectively, exhibited positive rho values, consistent with reduced responses to predictable words and increased responses with higher surprisal. We did not observe a difference in the magnitude of rho values between groups (**Figure 4C**).

**Figure 4.**
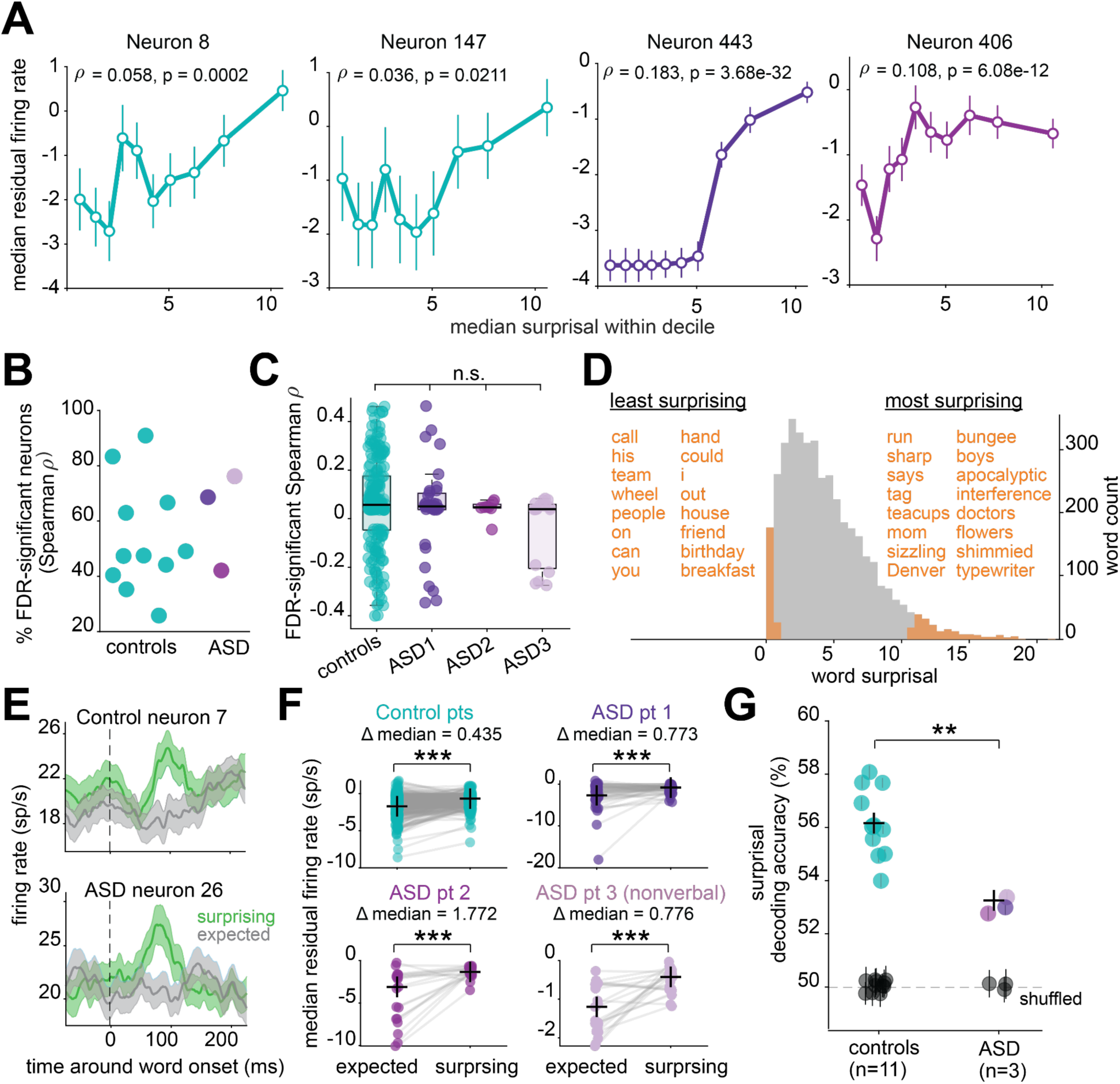
Population coding of word surprisal is altered in autism. **A,** Tuning curves for four example neurons showing residual firing rate as a function of word surprisal, binned into deciles. Points represent median firing rate within each bin and bars reflect IQR. **B,** Percentage of neurons per patient showing significant Spearman correlation with word surprisal after FDR correction. **C,** Spearman rho values for significant neurons; values were not different between patient groups (all P > 0.05, Monte Carlo subsampling test). **D,** Word surprisal values for all words in the data except the first word (n = 4,127). 205 words in orange reflect the top (surprising) and bottom (expected) 5% of surprisal values. **E,** Example peri-stimulus time histograms of firing rate responses to surprising or expected words from a hippocampal neuron in a control and autistic patient. **F,** For each neuron (circle), its word-averaged firing rates for surprising and expected words is plotted. Crosses reflect the median rate for each word type across neurons. Surprising words evoked significantly higher firing rates in each patient (P < 0.001, Wilcoxon signed-rank test on residual rates). **G,** For each patient, linear decoding accuracy (Support Vector Machine classifier) for surprising and expected words from neural population activity, for models using real and shuffled (black) data. Crosses reflect the group median; dashed line represents chance. Model accuracy was significantly reduced in autistic patients (P = 0.002, one-sided permutation test).

Because prediction error signals are expected to be strongest for highly unexpected inputs, we next examined neural responses at the extremes of the surprisal distribution. We identified the top and bottom 5% of words to categorize the most and least surprising words, respectively (**Figure 4D**). Examples of the most surprising words include, ‘bungee’, ‘apocalyptic’ and ‘typewriter’, while the least surprising words include, ‘call’, ‘hand’ and ‘birthday.’ Neural modulation to high- and low-surprisal words was directional: single hippocampal neurons exhibited higher firing rates to surprising words, a pattern observed across patient groups (**Figure 4E–F**; p<0.001, Wilcoxon signed-rank test).

Despite intact single-neuron prediction error signals, population-level decoding of surprising versus expected words was significantly reduced in each autistic patient compared to controls (**Figure 4G**; p < 0.01, one-sided permutation test), indicating that while individual neurons encode surprisal, this information is less effectively organized at the population level in autism. Although single neurons exhibit prediction-error sensitivity, increased shared variability across neurons (i.e., noise correlations) may increase redundancy and limit separability between surprising and expected words. This altered organization is consistent with our APG findings in autistic patients (**Figure 2E–K**), together suggesting atypical contextual integration rather than an absence of prediction error signaling.

### Reduced neural-linguistic feature alignment in autism

We next asked whether neural population activity reflects a shared structure across multiple linguistic dimensions, and whether this neural–linguistic alignment differs in autism. We used canonical correlation analysis (CCA), which identifies shared structure between neural population activity and a multidimensional linguistic feature space. For each patient, we applied cross-validated CCA to word-evoked firing rates across all neurons, the first 10 principal components of GPT-2 semantic embeddings, word surprisal, and polysemy (see **Methods**; results are consistent across different numbers of PCs and when using full embeddings). Along the first canonical component, neural activity and linguistic features were significantly correlated in all control patients and in ASD1, indicating a shared neural-linguistic representational axis (**Figure 5A**). ASD patients 2 and 3 exhibited markedly weaker correlations that did not reach significance (p < 0.001, permutation test; **Figure 5B**).

**Figure 5.**
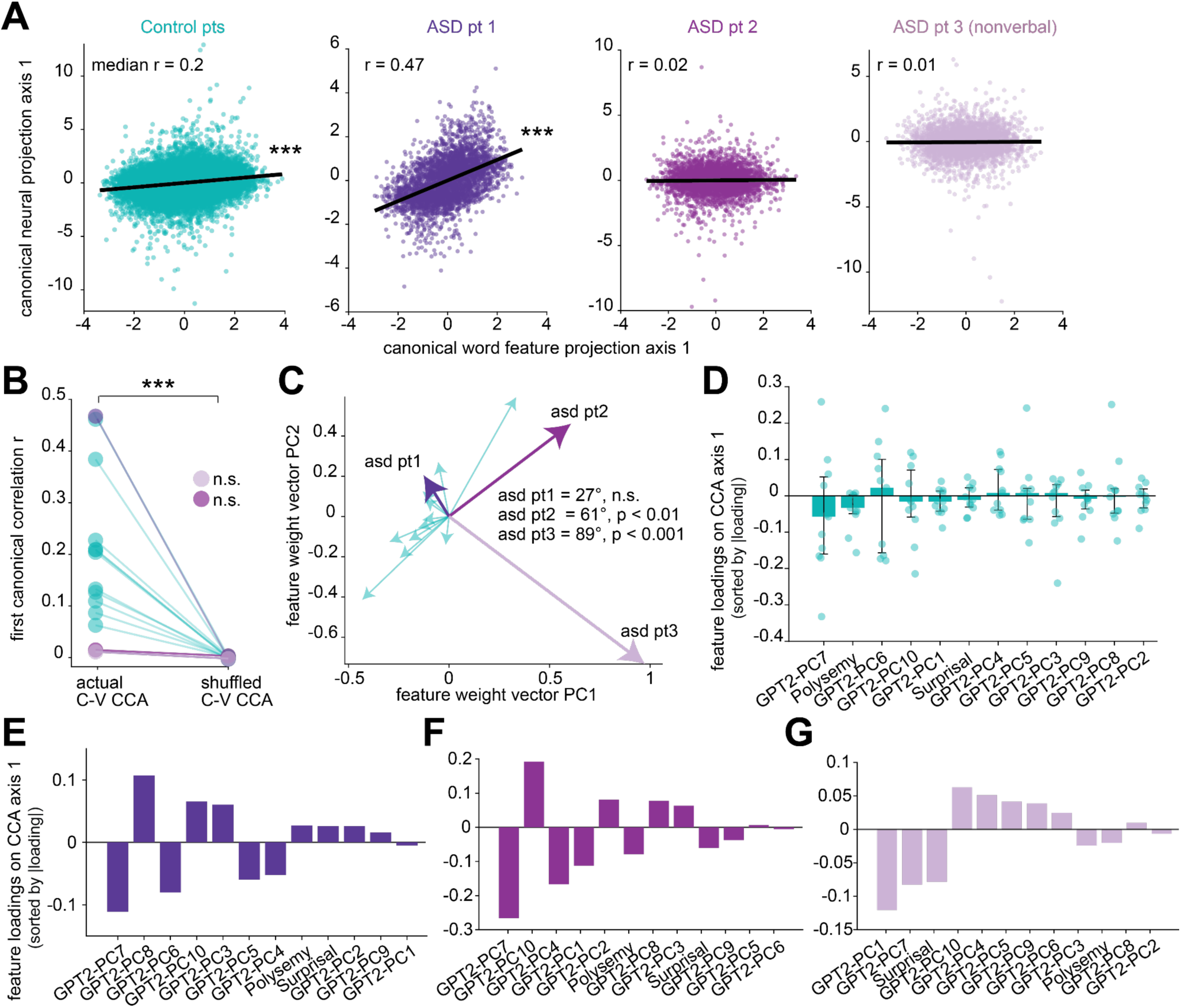
Absence of a shared linear neural-linguistic feature space in some individuals with autism. **A,** Each point represents a word, plotted by its held-out projection onto the first canonical axis of the linguistic feature space (n= 12 features) and the neural population space. Lines indicate linear fits. Control points are pooled across control patients. For each patient, the cross-validated axis-1 canonical correlation (r) quantifies the held-out association between neural and linguistic representations. Controls and ASD1 have significant correlations (P < 0.001, word-label permutation test). **B,** For each patient, the mean cross-validated canonical correlation on CCA axis 1 (actual) is shown alongside the median of the permutation null distribution, obtained by shuffling words of the linguistic feature matrix (P < 0.001, permutation test). **C,** For each patient, their canonical feature weight vectors for CCA axis 1 after sign alignment to the control mean and unit normalization. The plot shows the relative orientation of weight vectors in feature space (PCA visualization). For each ASD patient, cosine angle between their weight vector and the mean control vector is shown; significance was assessed using a bootstrap null derived from control-control angles. **D,** Bars show the median loading (Pearson correlation between each linguistic feature and the out-of-fold canonical score on axis 1) across controls for each feature, with error bars indicating IQR, sorted by absolute median loading. **E–G,** Individual autistic patients: bars show patient-specific feature loadings, sorted by absolute value. Loadings were sign-aligned to the control direction for comparability. Positive and negative values indicate the direction and strength of association between each feature and the canonical linguistic axis.

We compared the orientation of the canonical weight vector across patients after sign alignment and unit normalization (PCA projections are used only for visualization; **Figure 5C**). We quantified similarity between patients using the angle between unit weight vectors in the 13-dimensional feature space. ASD patients 2 and 3 showed larger angular deviations from controls (**Figure 5C**; p < 0.001, one-sided bootstrap test against control-derived null), whereas ASD1 fell within the expected range (**Figure 5C**; p = 0.79). This is notable because it indicates that, beyond differences in alignment strength captured by canonical correlations (**Figure 5A**), these ASD patients exhibit a different organization of linguistic feature weighting, suggesting heterogeneous alterations in how semantic and contextual information are integrated at the population level.

Next, we quantified which linguistic features exhibit the strongest loadings on the canonical axis. In control patients, semantic embeddings and polysemy exhibited the largest loadings, indicating that contextual semantic information primarily defined the shared neural–linguistic dimension (**Figure 5D**). In ASD1, semantic embeddings had the largest loadings, with word surprisal and polysemy contributing minimally to the canonical axis, suggesting reduced incorporation of contextual and predictive information into the population-level representation (**Figure 5E**). Loadings from ASD2 and ASD3 are shown for completeness; however, because a reliable shared axis was not identified in these patients, their loading patterns are not interpreted (**Figure 5F-G**). These results suggest that autism is not characterized by a uniform alteration of specific linguistic features but rather by variability in the existence and organization of a shared neural–linguistic subspace linking language features to population activity.

## DISCUSSION

We characterized the neural computations for semantic coding during language comprehension in three autistic patients. Despite heterogeneity in language ability, autistic patients exhibited a shared neural profile, characterized by intact core semantic representations but systematic alterations in semantic contextualization relative to controls. All autistic patients had reduced neural signatures of contextualization, including: (1) altered neural weighting of prior words for semantic coding a given word, (2) single neuron alignment to semantic embeddings from earlier, less contextual layers of GPT-2, (3) reduced effective dimensionality of the neural subspace predictive for semantics, and (4) reduced neural decoding of word surprisal. These findings align with models of altered predictive processing underlying perceptual deficits in autism (Pellicano and Burr, 2012; Brock, 2012; Mitchell and Ropar 2004; Lawson et al., 2014; Van de Cruys et al., 2014; Van Boxtel and Lu, 2013; Schneebeli et al., 2022; Noel and Angelaki, 2023). Relative to controls, they demonstrated comparable single-neuron semantic encoding, neural population discrimination between word meanings, and context-dependent tracking of word meaning. The quantitative similarity of these core semantic computations between control and autistic patients argues against the possibility that group differences arise solely from reduced attention or increased neural noise. Rather, our results suggest that the fundamental neural mechanisms supporting semantic representation during speech comprehension remain consistent in autism, while the integration of contextual information is selectively altered.

The broad distribution of neural APG values in autistic patients resembles the broader, less precise priors proposed in Bayesian theories of ASD (Pellicano and Burr, 2012; Mitchell and Ropar, 2004; **Figure 2I**). Broader priors imply that expectations are less constraining, such that perception depends more on incoming input than on context-based prediction. Thus, contextual information may still be used, but in a less focused and less selective manner. Even when the same previous word was identified as the strongest contextual contributor across patient groups, autistic patients’ neural weighting of that word was significantly altered - either exaggerated or attenuated - relative to controls. Together, these APG findings suggest altered scaling of contextual information during language processing.

It remains debated whether autistic perception is better characterized by reduced top-down influences on perception or, alternatively, enhanced bottom-up sensory-perceptual processes (Van de Cruys et al., 2014; Cannon et al., 2021; Van Boxtel and Lu, 2013; Brock 2012; Lawson et al., 2014; Pellicano and Burr, 2012). Our findings are more consistent with reduced top-down contextual influence. Single neuron phonemic and semantic encoding in autistic patients was comparable to controls. Additionally, single neurons in all autistic patients exhibited prediction error responses to surprising words. However, population-level decoding of surprisal was reduced, suggesting that prediction-related information is less effectively organized across the neural population. Lastly, neural responses in autistic patients were best predicted by embeddings from earlier GPT-2 layers that capture surface linguistic structure (e.g., word form and syntax), rather than deeper contextual semantic representations. Together, these findings suggest not an absence of prediction-error signaling in autism, but a change in how contextual predictions and their violations are distributed across neural populations. Overall, our findings are broadly consistent with Bayesian predictive coding accounts proposing altered weighting of prior expectations in autism, but they do not uniquely distinguish between competing formulations of these theories (Pellicano and Burr, 2012; Lawson et al., 2014; Van de Cruys et al., 2014; Chrysaitis and Series, 2023; Noel and Angelaki, 2023; Cannon et al., 2021).

Although single neurons in autistic patients aligned most strongly with earlier GPT-2 layers, population decoding still successfully reconstructed later contextual embeddings, indicating that contextual information is present in the neural population (**Figure 3E**). However, effective dimensionality of the neural subspace predictive for semantics was significantly lower in all autistic patients compared to controls. Thus, neural patterns used for predicting semantics are more redundant and less independent, which can limit the neural representational capacity for semantic coding. Said another way, in autistic patients, neural population activity is dominated by a smaller number of shared latent signals, causing neurons to vary more together rather than independently. This bias towards lower effective dimensionality in autistic patients could contribute to the reduced surprisal decoding and altered neural APG we observed, as lower effective dimensionality means there are fewer independent coding axes, which could impose less selective contextual constraint, leading to a broader distribution in the neural weighting of prior words.

In several analyses, semantic coding differed between autistic patients in how closely they resembled control patients. ASD patients 2 and 3 had lower language ability than controls, and their neural results often diverged from both control patients and ASD1, whose IQ and language ability were within the range of the control group. In contrast to controls and ASD1, ASD patients 2 and 3 showed higher firing rate responses to words, the most altered neural APG values - including negative correlations between neural and LLM-derived APGs - reduced population decoding performance for semantic embeddings, and an absence of a shared neural–linguistic feature space. These differences suggest that variability in language ability may be associated with differences in how semantic information is organized across neural populations.

More broadly, this work illustrates how large language models can serve as powerful computational tools for characterizing the neural basis of language and its alteration in neurological conditions. Large language models provided quantitative representations of contextual meaning and linguistic structure that allowed us to probe how different semantic features are organized within neural population activity and to gain insight into the neural mechanisms underlying language comprehension in autism. These results, therefore, argue for the applicability of new computational methods as a tool for uncovering the neurocomputational basis of psychiatric and neurological disease.

## METHODS

### Patient demographics

All experiments were performed in the Epilepsy Monitoring Unit (EMU) at Baylor St. Luke’s Hospital using standardized approaches (Xiao et al., 2024A and B). Experimental data were recorded from 14 patients with medically refractory epilepsy undergoing clinical standard-of-care intracranial seizure monitoring, 3 of whom had comorbid ASD, and 11 of whom who did not (**Extended Data Fig. 1**). ASD patients were 22, 19, and 34 years old, respectively. The remaining patients were allistic, and collectively constitute our control group. The control patients were matched for intractable epilepsy status and had an age range of 18-48 (mean 30). ASD patient 1 (ASD1) was autistic without intellectual disability (IQ, verbal, and perceptual scores fell within the control range - 30th, 20th, and 30th percentile respectively; Wechsler Adult Intelligence Scale IV). ASD patient 2 (ASD2) was autistic with mild intellectual disability (although their IQ score of 62 fell within the control range, 20th percentile) and impaired verbal comprehension (VCI = 68, lower than all controls; Perceptual reasoning (PRI) fell within control range - 20th percentile; Wechsler Adult Intelligence Scale IV). ASD patient 3 (ASD3) had profound autism (nonverbal with severe Intellectual Disability; functioning level 12 months; Adaptive Behavior Composite score 30 - Vineland Adaptive Behavior Scale).

### Human intracranial neurophysiology

Single neuron data were recorded from stereo-electroencephalography (sEEG) electrodes with microwires extending from the tip (AdTech Medical Behnke-Fried). Each patient had an average of 3 probes terminating in the left and right hippocampus. Electrode locations were verified by co-registered pre-operative MRI and post-operative CT scans. Each probe includes 8 microwires, each with 1 contact, specifically designed for recording single-neuron activity. Single neuron data were recorded using a 512-channel *Blackrock Microsystems Neuroport* system sampled at 30 kHz. To identify single neuron action potentials, the raw traces were spike sorted using the *Wave_clus* sorting algorithm (Chaure et al., 2018) and then manually evaluated. Noise was removed and each signal was classified as multi or single unit using several criteria: consistent spike waveforms, waveform shape (slope, amplitude, trough-to-peak), and exponentially decaying ISI histogram with no ISI shorter than the refractory period (1 ms). The analyses here used all single and multiunit activity (see **Extended Data Fig. 2**). The number of neurons per patient in the control group ranged from 11-54; the number of recorded neurons in ASD patients was 51, 19, and 21, respectively.

### Electrode visualization

Electrodes were localized using the software pipeline intracranial Electrode Visualization (iELVis; Groppe et al., 2017) and plotted across patients on an average brain using Reproducible Analysis & Visualization of iEEG (RAVE; Magnotti et al., 2020). For each patient, DICOM images of the preoperative T1 anatomical MRI and the postoperative Stealth CT scans were acquired and converted to NIfTI format (Li et al., 2016). The CT was aligned to MRI space using FSL (Jenkinson et al., 2002). The resulting coregistered CT was loaded into BioImage Suite (version 3.5β1; Joshi et al., 2011) and the electrode contacts were manually localized. Electrodes coordinates were converted to patient native space using iELVis MATLAB functions (Yang et al., 2012) and plotted on the Freesurfer (version 7.4.1) reconstructed brain surface (Dale et al., 1999). Microelectrode coordinates are taken from the first (deepest) macro contact on the Ad-Tech Behnke Fried depth electrodes. RAVE (Magnotti et al., 2020) was used to transform each patient’s brain and electrode coordinates into MNI152 average space. The average coordinates were plotted together on a glass brain with the hippocampus segmentation and colored by patient.

### Natural language stimuli

Patients with epilepsy listened to three episodes (5- to 13-minutes in duration) taken from *The Moth Radio Hour,* totaling 27 minutes of listening time (4,128 words). The three stories were, “Life Flight”, “The Tiniest Bouquet”, and “The One Club”. In each story, a single speaker tells an autobiographical narrative in front of a live audience. The three selected stories were chosen to be both interesting and linguistically rich. Stories were played continuously through the built-in audio speakers of the patient’s hospital television. The audio signal was synchronized to the neural recording system via analog input going directly from the computer playing the audio into the Neural Signal Processor at 30 kHz.

### Audio Transcription

After experiments, the audio .wav file was automatically transcribed using Python and Assembly AI, a state-of-the-art AI model to transcribe and understand speech. The transcribed words and corresponding timestamp output from Assembly AI was converted to a TextGrid and then loaded into Praat, a software for speech analysis. The original .wav file was also loaded into Praat and we manually checked the spectrograms and timestamps, correcting each word to ensure the word onset and offset times are accurate. The TextGrid output of corrected words and timestamps from Praat was converted to a .xls and loaded into Matlab and Python for further analysis.

### Semantic embedding extraction from language models

**1) Word2Vec.** We used the pre-trained *fastText Word2Vec* model in MATLAB to extract word embeddings for all words in our dataset (Joulin et al., 2016; Mikolov et al., 2013). This pre-trained model provides 300-dimensional word embedding vectors, trained on millions of words text, to capture semantic relationships between words. Any surname words, such as “Harwood” or proper nouns like “Applebee’s” that did not have word embeddings were discarded from the analysis (these were rare in our sample).
**2) GPT-2 Large**. In an excel file of our transcribed words, each row corresponded to an individual token, including punctuation and sentence terminators such as “.”, “?”, and “!”. We reconstructed sentences by iterating through the file and appending words until encountering a recognized sentence delimiter. This process yielded multi-word segments that more accurately captured natural linguistic boundaries. Any remaining words following the last delimiter were grouped into a final sentence. Notably, punctuation tokens were preserved to maintain contextual fidelity but were tracked carefully to avoid introducing sub-word alignment errors during tokenization.

To extract GPT-2 embeddings, we employed the “gpt2-large” model (Radford et al., 2019) from the Hugging Face Transformers library (Wolf et al., 2020). This version of GPT-2 features a total of 37 layers (1 initial embedding layer plus 36 transformer layers), each producing 1280-dimensional hidden states, with a maximum context length of 1024 tokens. Although GPT-2 supports a 1024-token context window, all embeddings used in this study were computed within individual sentences only - that is, the model’s context was reset whenever a sentence-ending delimiter (e.g., “.”, “!”, “?”) was encountered. We extracted hidden states from all 37 layers using a progressive context approach designed to incorporate GPT-2’s autoregressive nature. Specifically, for each sentence, we began with an empty context and added words sequentially. After adding each new word, we tokenized the updated text string via GPT-2’s Byte Pair Encoding (BPE) tokenizer, ran the token sequence through the model in no-grad (inference-only) mode, and retrieved the hidden states. We then identified the newly appended word’s sub-word tokens and averaged their corresponding vectors to obtain a single word-level embedding. Because GPT-2 often splits words into multiple sub-word fragments (otherwise known as “tokens,” e.g., “computa” + “tion”), averaging those fragments preserves consistency with the transcript’s original word boundaries. The GPT-2 outputs resulted in 1280-dimensional embeddings for each layer.

### Firing rate responses to words

We computed the duration for each word in the story and for each non-word in Jabberwocky stimuli. The firing rate of a neuron for each word was the number of spikes occurring during the word with an 80 ms delay after word onset to account for the approximate delay of information to hippocampus. This value was divided by word duration and then multiplied by 1000 to compute spikes per second.

### Word Surprisal

We quantified how predictable each word in the story was to a large language model according to previously described methods (Gwilliams et al., 2024; Weissbart and Martin, 2024). For each word, w_i_, we maintained a progressive left-to-right transcript and evaluated its predictability under a series of context windows. Specifically, we extracted the last C ∈ {4,8,16,32,64,128,256,512,1024} tokens preceding *w_i_*, fed this context into GPT-2 Large, and obtained the model’s predicted probability distribution over the vocabulary for the upcoming word (**Figure 4D**):

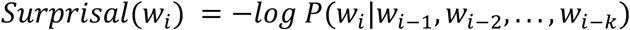

### Polysemy

To compute polysemy for each word, we computed cosine distance between each GPT-2 layer 25 embedding vector of all occurrences of a given word (at least 2 occurrences, 396 words total). We took the median of this distribution to obtain one polysemy value per word. We repeated this analysis using neural embeddings instead of GPT-2 embeddings to compute the neural cosine distance values, or neural polysemy. We also computed polysemy for each word as the total number of synsets (i.e., distinct senses) associated with each word in WordNet (Fellbaum, 1998). These polysemy values yielded the same results shown in **Figure 2D**.

### Regressing spike counts on word embeddings, phonemes, and linguistic features

Word embeddings, such as those derived from Word2Vec or similar models, are typically high-dimensional and can exhibit substantial collinearity among dimensions - that is, many of the embedding features are correlated with one another. We first used Principal Component Analysis (`pca`, Matlab 2023b) on the full word embeddings to obtain uncorrelated features that still capture the dominant structure in the embedding space with reduced dimensionality. For each word, we used the first 100 principal components (PC) from Word2Vec or GPT-2 embeddings because they explained at least 60% of the variance of the original embedding vectors.

We modeled the spike count responses of individual neurons using a Poisson Generalized Linear Model (GLM) with a log-link function and ridge (L2) regularization (we confirmed that spike counts for each word follow a Poisson distribution; **Figure 1G and Figure 3A-B**). The model aimed to predict the number of spikes a neuron fired in response to each word (summed spikes across word duration), using the top 100 principal components (PCs) of the word embedding vector, the duration of the word, and the interaction between each PC and the word’s duration as predictors (z-scored prior to model fitting). This resulted in a 201-dimensional feature vector for each word: 100 PC values, 1 duration value, and 100 PC×duration interaction terms.

The Poisson GLM assumes that the spike count *y_i_* for word *i* is drawn from a Poisson distribution with mean λᵢ where:

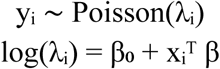

Where 𝜆*_i,n_*, the expected spike count for word (*i*) and neuron n (*n*), is modeled by the linear regression coefficients (*β*), one for each of the 201 predictors (x_i_^T^), plus the y-intercept (*β*_0_).

For each neuron, we fit a Poisson ridge regression model using a nested cross-validation framework with 5 contiguous outer folds. Words were partitioned into five sequential test blocks of approximately equal size, such that in each outer fold one contiguous block of words served as the held-out test set and the remaining four blocks were used for training. This blocked outer-fold design was used to reduce temporal leakage arising from autocorrelation across nearby words in the continuous speech stream. Within each outer training set, the optimal regularization strength (α; alpha) was selected using 5-fold inner cross-validation, drawn from a log-spaced range (10^-3^ to 10^3^), and the value maximizing the mean validation log-likelihood across inner folds was selected. Using the selected α, a Poisson ridge model was then fit to the full outer training set and then evaluated on the held-out test set using several performance metrics, including log-likelihood and adjusted R². To assess model complexity, we estimated the effective degrees of freedom (EDF) from the training data and used this to compute an out-of-sample AIC-like metric for the test set (**Figure 1J**). Final summary statistics for each neuron were obtained by taking the median across the 5 outer folds.

To assess statistical significance relative to a null, we generated 100 shuffled predictor matrices per dataset, in which embeddings were shuffled while keeping word duration preserved. Each shuffle circularly shifted the embedding matrix along the observation axis by a randomly selected offset of at least ±100 samples (to avoid short-range autocorrelation artifacts). To determine if the actual model showed significant improvement from the shuffled, we performed a permutation test comparing the actual model LLH to the shuffled model LLH distribution. We then took the difference between the actual and shuffled model (mean) LLH (LLH diff; ΔLLH). A neuron was fit by the model if its ΔLLH was both positive and significant and if McFadden’s pseudo R² value was positive (P < 0.05, permutation test, **Figure 1H**).

Predictor significance was assessed using p-values from a Poisson generalized linear model fit to the same predictors, while coefficient magnitudes were taken from the L2-regularized Poisson regression model used for prediction. As expected, word duration was always a significant predictor, followed by the first eight embedding dimensions and their interactions with duration, with the number of significant predictors gradually decreasing across higher dimensions.

#### Phonemic regression

We built a fixed-length feature space over the 43 IPA phonemes observed in our corpus. Here, IPA refers to the International Phonetic Alphabet, a standardized set of symbols that denote speech sounds independent of spelling. The podcast transcript was processed word by word: tokens were lower-cased and converted from graphemes to phonemes using the Python phonemizer package (version 3.0.1) with the eSpeak backend as the grapheme-to-phoneme (G2P) model (https://pypi.org/project/phonemizer/3.0.1/). For each word, we took the canonical IPA transcription returned by phonemizer. Phonemes were then manually inspected and corrected by linguists (JB and HZ) in Praat.

Each word was then represented as a 43-dimensional count vector whose entries reflect the number of occurrences of each IPA phoneme in that word (unigram counts; no duration or stress information). This approach is similar to Khanna et al., in that words are represented by the presence of their constituent phonemes; however, we did not group phonemes into broader classes, in order to preserve the full identity of the 43 phones observed in our data and maintain the embedding’s informational richness and dimensionality (Khanna et al., 2024). To better differentiate words that contain the same phoneme set but in different orders (e.g., dog vs god), we double-counted the first phoneme in every word while counting all subsequent phonemes once. The resulting word-by-phoneme matrix (N words × 43 features) was used in subsequent analyses relating phonetic composition to neural responses. We used this word-by-phoneme matrix, word duration, and their interactions in the same Poisson regression described above for predicting spike counts of the current word. The data folds used for training and testing were preserved across semantic and phonetic models and both models had the same number of words.

We used the same Poisson ridge regression model to predict an individual neuron’s spike count for each word from 61 linguistic features: (1) full phonetic embeddings (43 IPA phones; count coded, cf. Khanna et al., 2024), (2) 10 PCs of semantic embeddings, (3) word order in sentence, (4) word order in clause (Stanford CoreNLP; Manning et al., 2014), (5) opening node number (Stanford CoreNLP; Manning et al., 2014), (6) closing node number (Stanford CoreNLP; Manning et al., 2014), (7) word frequency (SUBTLEX-US; Brysbaert & New, 2009), (8) pitch, (9) a scalar representing degree of polysemy (Cruse et al., 1986; Landauer, 2001; Ethayarajh, 2019) and (10) word duration (**Extended Data Figs. 3-4**).

### Embedding distance

The semantic embedding vector for each word was its full vector of embedding values extracted from GPT-2 layer 25 (1280 in length) language model. For each patient, for each word, its neural embedding vector was every neuron’s firing rate response for that word (11-54 neurons in length). We used the function `pdist`, Matlab 2023b, to compute cosine distance between semantic embedding vectors of all 8,514,001 word pairs, known as semantic distance. We repeated the procedure, computing cosine distance between neural embedding vectors for all word pairs, termed neural cosine distance. “All word pairs” included pairs of the same word. We correlated semantic distance and neural distance from all word pairs to compute the correlation coefficients in **Figure 2B**.

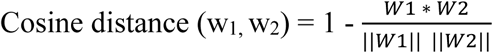

Cosine distance between neural or semantic embeddings of word 1 and word 2 is the 1 minus the dot product of embedding vectors for each word, divided by the magnitudes (norms) of each word vector.

### Attention Pattern Grid

The ***empirical attention pattern grid*** is an estimate of the degree to which all past words influence each current word in terms of the proportion of influence they have on the semantics of that word (Katlowitz et al., 2025). It therefore recapitulates the function performed by the APG used in transformer architecture LLMs. We first estimated the neural APG from neural data alone and, for comparison, performed an analogous process on the LLM embeddings. Note that the LLM-derived empirical APGs were not derived directly from the LLM APGs themselves; they were inferred so as to keep the estimation procedure the same for brain and LLM. For reference, a true APG from an LLM is the result of the dot product of the Query and Key matrices, and there are many heads of attention per layer.

Our empirical APGs were calculated in the same manner for all embedding matrices.

Embedding matrices were 4,128 rows, corresponding to the number of words in our stimulus set, by M columns, corresponding to the number of embedding dimensions per word. For the neural embedding matrix M was equal to the number of neurons per patient (mean 30). For the semantic empirical APGs, M was 1,280 (GPT-2, all layers; **Figure 2E-K**).

We then created two new embedding vectors for each word: the average vector across all instances of that word 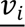 and the shifted representation of the word j is 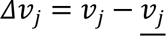. Then each value of the APG was created by measuring the cosine similarity between the shifted vector and the mean vector of the previous words

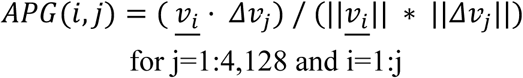

Words with only one instance in the entire task were excluded from analysis as it was impossible to calculate a shifted representation. Only causal influences were calculated.

### Reduced Rank Regression

To quantify how hippocampal population activity predicts high-dimensional semantic embeddings of heard words, we applied Reduced Rank Regression (RRR) using the Matlab code and following the framework of Semedo et al. 2019. For each patient, neurons served as predictors and individual words as observations (**Figure 3C**).

Neural data preprocessing: For each patient, we computed per word firing rates normalized by word duration (see *Firing rate responses to words).* The resulting matrix contained one observation per word and one column per neuron (X). Word-level confounding variables - the log of word frequency and position - were regressed out of the neural activity matrix, and the residuals were used in analysis.

The first 30 Principal Components of semantic embeddings from each layer of GPT-2 were used in the regression, forming a word by semantic embedding matrix (Y). So, a model was fit separately for each GPT-2 layer and patient. RRR estimates a mapping from neural activity X to semantic embeddings Y that is constrained to lie in a low-dimensional shared subspace. We used the Semedo et al., 2019, `ReducedRankRegress` function, which implements the closed-form solution for:

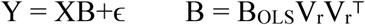

where V_r_ are the top r eigenvectors defining the predictive subspace of the OLS mapping. The regression coefficient matrix B is constrained to rank r. For each patient, this decomposition yields:

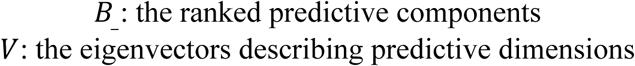

These define the semantic subspace captured by the neural population. To determine the number of predictive dimensions, we evaluated models with 0–15 RRR dimensions using 10-fold cross-validation, following Semedo et al. Model performance (averages across folds) was quantified as the cross-validated variance explained (1 – NSE) across predictive subspace dimensionalities (ranks = 0 - 15). The optimal dimensionality (r; rank) can be defined as the number of dimensions needed for the prediction accuracy to plateau, selected using the Semedo et al., 2019 function, `ModelSelec`t. These RRR results quantify how many dimensions of semantic variation can be linearly predicted from hippocampal population activity, as well as the geometry of the neural to semantic mapping.

#### Effective dimensionality

We quantified the effective dimensionality of the semantic-predictive subspace using the participation ratio of the singular-value spectrum of the top r predictive dimensions B_(:,1:r). Effective dimensionality measures how evenly predictive variance is spread across the latent dimensions (ranks). Using each patient’s optimal rank or the median optimal rank of control patients when computing effective dimensionality at each layer yielded the same results (**Figure 3G**).

### Support Vector Machine Decoder

We used a SVM decoder (Bishop, 2006) with a linear kernel to determine whether firing rates during words carry information about word surprisal by distinguishing the least and most surprising words (**Figure 4G**). For each patient, we used the firing rate responses from every neuron (n = 11-54 neurons per patient) for the most surprising (class 1) and least surprising (class 2) words to predict the class for each word from these neural data. There were 204 words in each class. To train and test the model, we used a tenfold cross-validation. In brief, the data were split into ten subsets, and in each iteration the training consisted of a different 90% subset of the data; the testing was done with the remaining 10% of the data. We used the default hyperparameters as defined in `fitcsvm`, MATLAB 2023b, and z-scored normalization of firing rates. Decoder performance was calculated as the percentage of correctly classified test trials. We compared model performance for predicting train and test data to check for overfitting. In each iteration, we trained a separate decoder with randomly shuffled class labels. The performance of the shuffled decoder was used as a null hypothesis for the statistical test of decoder performance.

### Canonical Correlation Analysis

To quantify the shared linear structure between neural population activity and linguistic features, we performed canonical correlation analysis (CCA) using the `canoncorr` function in Matlab 2023b. For each patient, neural data consisted of a word-by-neuron firing rate matrix X (words × neurons). Linguistic features were represented as a word-by-feature matrix Y (words × 12 features), including a minimum 10 principal components of GPT-2 semantic embeddings, word surprisal, and polysemy. Matrices X and Y were z-scored. CCA finds linear projections of the neural and linguistic feature spaces that maximize their correlation. Specifically, CCA identifies projection matrices *A* and *B* such that:

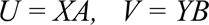

where *U* and *V* are the neural and linguistic canonical scores, respectively. The first pair of canonical variables corresponds to the projections that maximize the correlation between *U* and *V*. We implemented a 10-fold outer cross-validation procedure to estimate out-of-sample canonical correlations. Words were partitioned into 10 contiguous folds. For each fold, CCA weights were learned on the training set and neural and linguistic data from the held-out fold were projected onto the first pair of canonical axes. We then computed the Pearson correlation between held-out neural and linguistic canonical scores (r1). To assess whether observed canonical correlations exceeded chance levels, we constructed a permutation-based null distribution. For each of 1,000 permutations, word labels of the linguistic feature matrix were randomly shuffled and the identical 10-fold cross-validation procedure was repeated using the permuted feature matrix. Then we computed the mean cross-validated canonical correlation for the shuffled model, rPerm. For each patient, we computed the permutation p-value as the proportion of rPerm values greater than or equal to the observed cross-validated r1 (**Figure 5A-B**). We then fit CCA to the full dataset to obtain canonical weights and loadings for interpretation.

#### Comparison of canonical feature-weight vectors across patients

From the model fit on the full data, we obtained the feature-side canonical weight vector for the first canonical component. Because canonical axes are defined up to a sign flip, we performed sign alignment prior to comparing weights across patients. Specifically, we selected a seed control patient and flipped the sign of each control weight vector when its dot product with the seed was negative. We then computed the mean control direction from these aligned control vectors and used this mean direction as a reference to sign-align all patients (controls and ASD) by flipping any vector with a negative dot product relative to the reference. To compare feature-weighting patterns independent of scale, aligned weight vectors were normalized to unit length. Similarity between patients was quantified using the angular distance between unit vectors, expressed in degrees.

Significance of ASD deviations was assessed using a bootstrap null distribution constructed by resampling control patients and computing typical control–control angular deviations; p-values reflect the probability that the observed ASD angle exceeded this null distribution (**Figure 5C**).

#### Canonical structure loadings and sign alignment

For each linguistic feature *j*, the structure loading was defined as the Pearson correlation between the original feature time series and the out-of-fold canonical variate on the linguistic side (V₁ᴼᴼᶠ), computed across word tokens:

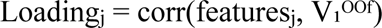

Because canonical variates in CCA are defined only up to an arbitrary sign, loading vectors were sign-aligned to enable meaningful comparisons across patients. First, loading vectors for control patients were aligned to a common orientation by flipping the sign of any individual control loading vector whose dot product with the control-group mean loading vector was negative. The resulting aligned control loadings were then used to define a control reference direction. Loading vectors for autistic pateints were subsequently sign-aligned to this control reference direction using the same dot-product criterion. This procedure preserves the relative pattern and magnitude of feature associations while removing arbitrary sign flips inherent to CCA. For control patients, group-level summaries were computed using the median structure loading across patients for each feature, with uncertainty estimated by bootstrap resampling across control patients (**Figure 5D-G**).

### Statistics

To assess statistical significance while accounting for differences in neuron counts and small patient sample sizes, we used nonparametric randomization and permutation tests appropriate to the structure of each analysis. For single-neuron analyses (**Figure 1**), significance between control and autistic patients was assessed using a randomization test. For each autistic patient, the observed statistic (e.g., median firing rate or median LLH) was compared to a null distribution generated from 5,000 neuron-count–matched subsamples of control neurons (sampling without replacement within each subsample). P-values were computed as the fraction of null medians at least as extreme as the observed value.

Permutation tests (**Figure 3**): To assess whether a value differed between autistic and control patients, we performed a nonparametric permutation test at the patient level. For each autistic patient, we first computed the observed test statistic as the median across GPT-2 layers of the difference between that patient’s value and the control group median at each corresponding layer. To construct a null distribution reflecting typical between-subject variability among controls, we repeatedly (10,000 iterations) selected one control patient at random to serve as a “pseudo-ASD” subject, with the remaining controls forming the reference group. For each iteration, we computed the same median difference statistic between the pseudo-ASD patient and the median of the remaining controls. This procedure generated a null distribution of control-versus-control differences under the assumption of no group effect. Statistical significance was assessed using a one-sided test, quantifying the proportion of null statistics less than or equal to the observed statistic (testing whether autistic patients exhibited lower values than controls).

Decoding accuracy (**Figure 4G**): We performed a nonparametric permutation test on group mean accuracy. Under the null hypothesis that group labels were exchangeable, patient labels were randomly permuted (10,000 iterations), assigning three patients to the ASD group and the remaining patients to the control group on each iteration. For each permutation, the test statistic was computed as the difference in mean accuracy between the ASD and control groups. The observed group difference was compared to the resulting null distribution to obtain a one-sided p-value, testing the hypothesis that decoding accuracy was different (lower in surprisal) in ASD than in controls.

## Acknowledgements

We thank Joshua Adkinson and Victoria Gates for invaluable assistance.

## Extended Data Figures

**Extended Data Fig. 1.**
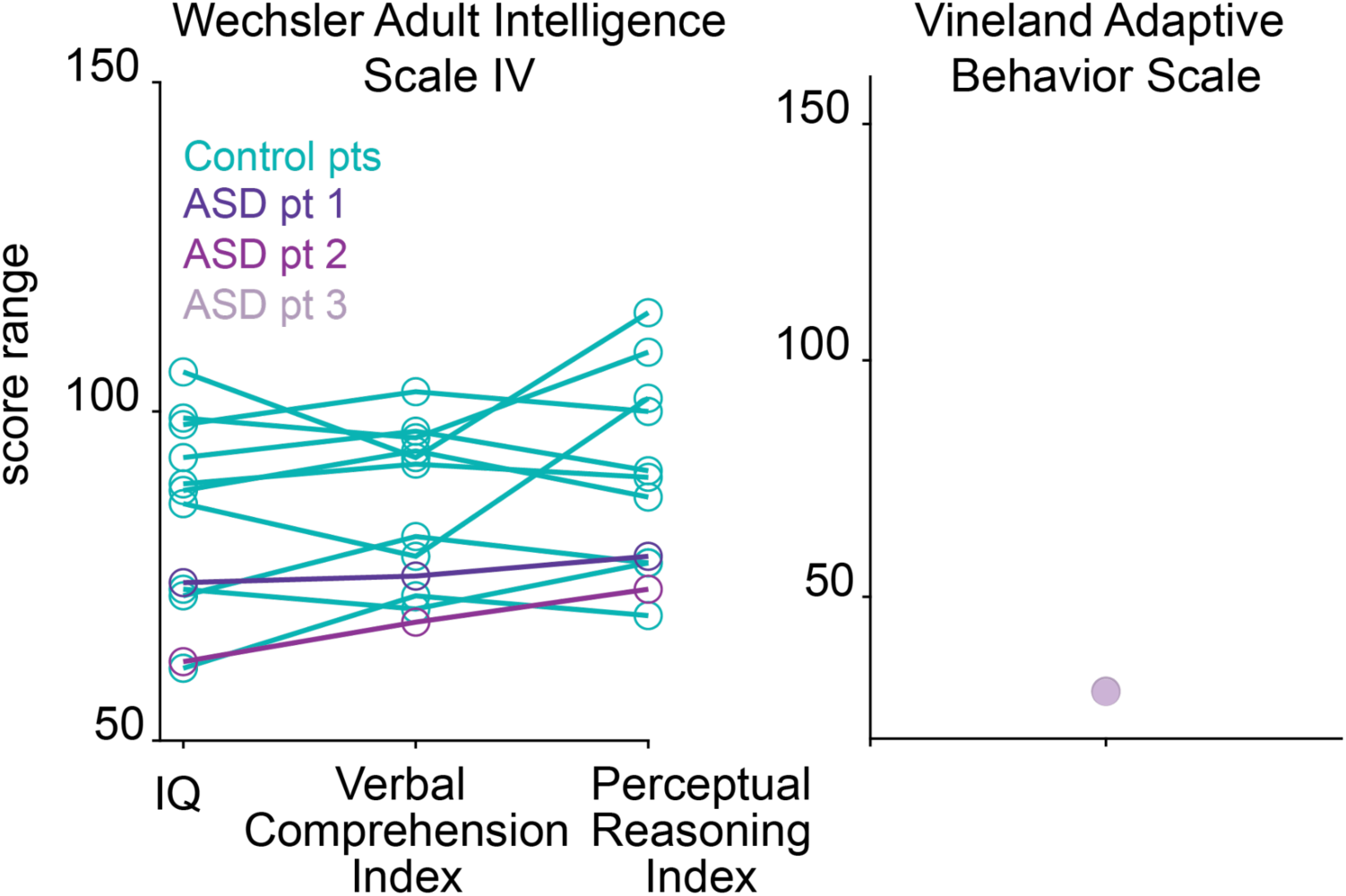
Test scores from 10 control patients and each ASD patient for cognitive and clinical assessments that were conducted. Assessments were not completed for one control patient.

**Extended Data Fig. 2.**
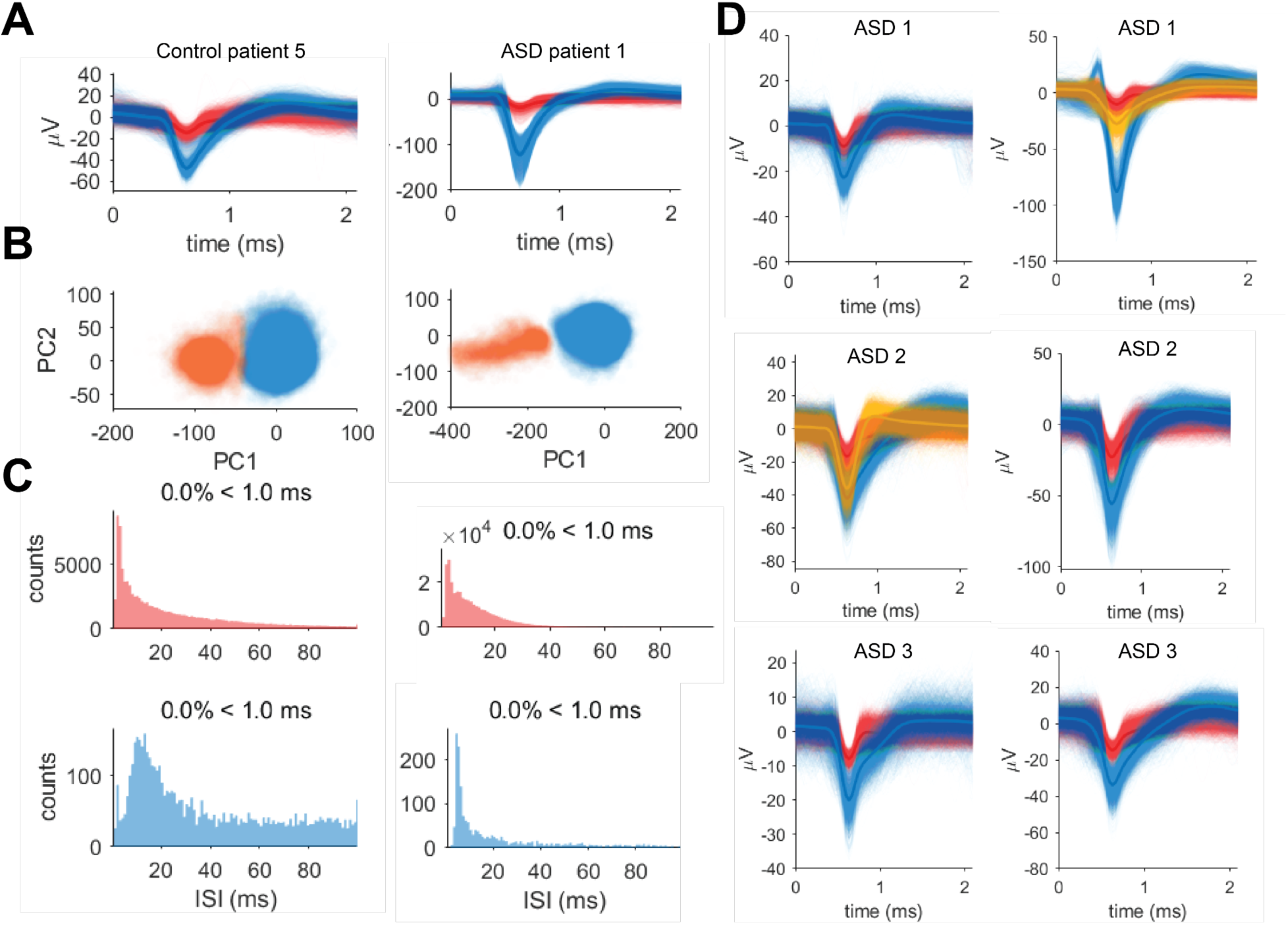
Single and multi-unit detection and isolation. **A,** A random sample of 5,000 individual waveforms from a single hippocampal neuron (blue) and multi unit cluster (red) from control patient 5 and ASD1 recorded on one electrode channel. Bolded lines represent average waveforms. Noisy signals were already removed using *Wave_clus* sorting algorithms and manual inspection. **B,** For each patient, all waveforms from each cluster in A are well isolated in Principal Component space. **C,** Interspike interval (ISI) histograms from the neurons in A. Single and multi-unit clusters always have refractory period violations below 5% (as shown in figure titles). **D,** Example hippocampal single (yellow and blue) and multiunit (red) waveforms on individual electrode channels from three autistic patients that were identified and sorted following the criteria in B-C using *Wave_clus* software and manual inspection.

**Extended Data Fig. 3.**
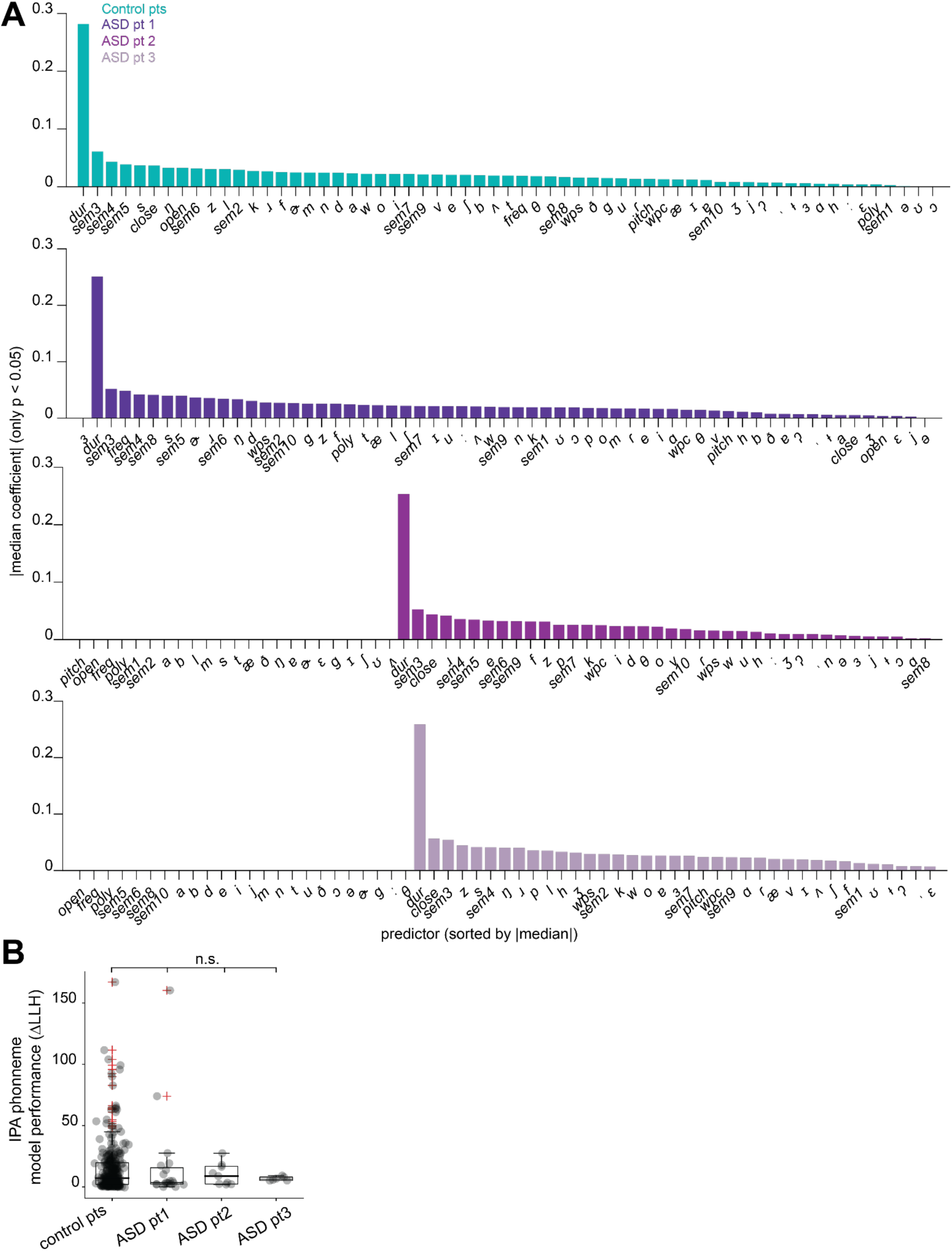
Phonemic and combined linguistic feature regression models. **A,** The absolute value of the coefficient, or weight, of each predictor in the model is shown on the y-axis. Bars reflect the median predictor weight taken across neurons fit by the model. Predictors are sorted by decreasing weights. Poly = polysemy; Wps = word position in sentence; wpc = word position in clause; freq = word frequency; open and close represent opening and closing nodes; sem1-10 are semantic embeddings from GPT-2 layer 25. Some predictors were not significant for any neurons in ASD patients 2-3. **B,** Log-likelihood improvements (ΔLLH; model performance) for fitted neurons from Poisson ridge regression predicting neuronal spike counts from word-level IPA phoneme embeddings using the equation in Figure 1G. Autistic patient performance is comparable to controls (P = 0.07, 0.62. and 0.42 for ASD1-3 respectively, one-sided Monte Carlo subsampling test on medians using 5,000 neuron-matched resamples from controls).

**Extended Data Fig. 4.**
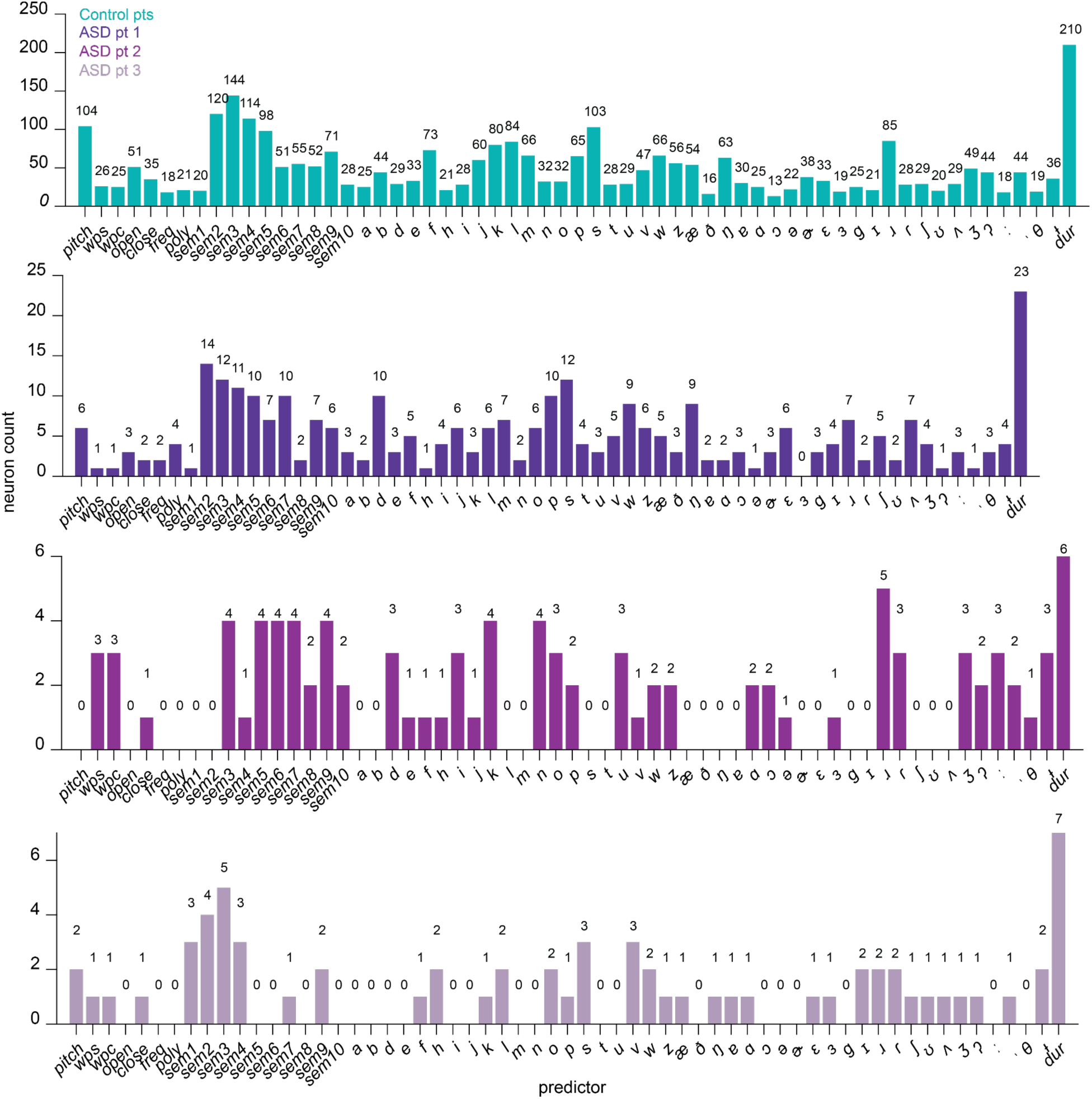
Regressing neuronal activity with multiple linguistic features. The number of neurons for which each predictor was significant in the model is plotted on the y-axis, and the total number of neurons is also displayed on top of the bar plot. This model used semantic embeddings from GPT-2 layer 25. Predictors and values correspond to those in Extended Data Fig. 3A. Some predictors were not significant for any neurons in ASD patients 2-3.

**Extended Data Fig. 5.**
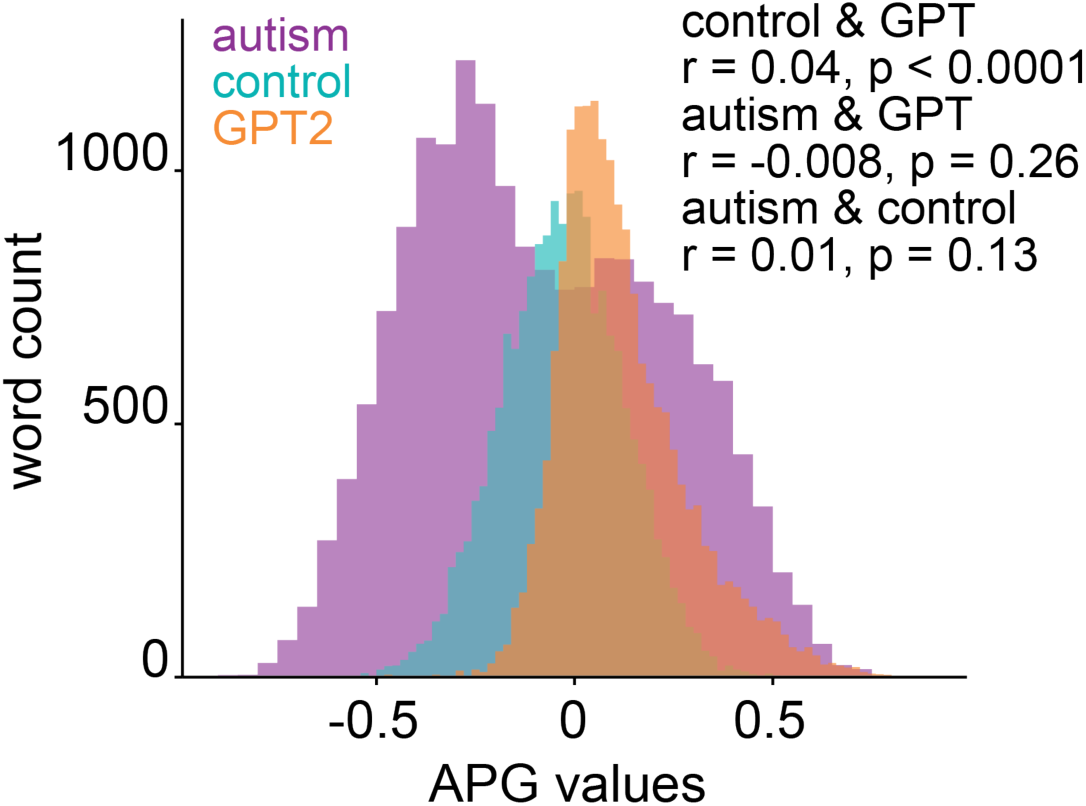
Neural APG distribution from ASD patient subset. The distribution of APG values taken 5 words before each word from the LLM (GPT2-Layer 26 embeddings), control patients, and ASD patients 2-3. Median APG values were taken across patients in each patient group. Pearson correlations between APG values in each group are shown.

## REFERENCES

Adams, R. A., Huys, Q. J., & Roiser, J. P. (2016). Computational psychiatry: towards a mathematically informed understanding of mental illness. Journal of Neurology, Neurosurgery & Psychiatry, 87(1), 53–63.

Alamia, A., Schwenk, J. C., Wagemans, J., & Sapey-Triomphe, L. A. (2026). Oscillatory traveling waves during visual entrainment in autistic and neurotypical adults. bioRxiv, 2026-01.

American Psychiatric Association. (2013). Diagnostic and statistical manual of mental disorders (5th ed.).

Aylward, E. H., Minshew, N. J., Goldstein, G., Honeycutt, N. A., Augustine, A. M., Yates, K. O., … & Pearlson, G. D. (1999). MRI volumes of amygdala and hippocampus in non–mentally retarded autistic adolescents and adults. Neurology, 53(9), 2145–2145.

Banker, S. M., Gu, X., Schiller, D., & Foss-Feig, J. H. (2021). Hippocampal contributions to social and cognitive deficits in autism spectrum disorder. Trends in neurosciences, 44(10), 793–807.

Bhamidimarri, P. M., Alhosani, K., Cai, H., Al-Ali, H., Abukhaled, Y. M., Tawamie, H., … & Hamdan, H. (2025). Review on the role of hippocampus in autism spectrum disorder: Recent insights into neuropathology, genetics, and emerging therapeutic strategies. Neurobiology of Disease, 107227.

Binder, J. R., Desai, R. H., Graves, W. W., & Conant, L. L. (2009). Where is the semantic system? A critical review and meta-analysis of 120 functional neuroimaging studies. Cerebral cortex, 19(12), 2767–2796.

Bishop, C. M. (2006). Pattern recognition and machine learning. New York : Springer

Blanchard, T. C., Wolfe, L. S., Vlaev, I., Winston, J. S., & Hayden, B. Y. (2014). Biases in preferences for sequences of outcomes in monkeys. Cognition, 130(3), 289–299.

Brock, J. (2012). Alternative Bayesian accounts of autistic perception: comment on Pellicano and Burr. Trends in cognitive sciences, 16(12), 573–574.

Brodski-Guerniero, A., Naumer, M. J., Moliadze, V., Chan, J., Althen, H., Ferreira-Santos, F., … & Wibral, M. (2018). Predictable information in neural signals during resting state is reduced in autism spectrum disorder. Human brain mapping, 39(8), 3227–3240.

Brysbaert, M., & New, B. (2009). Moving beyond Kučera and Francis: A critical evaluation of current word frequency norms and the introduction of a new and improved word frequency measure for American English. Behavior research methods, 41(4), 977–990.

Cannon, J., O’Brien, A. M., Bungert, L., & Sinha, P. (2021). Prediction in autism spectrum disorder: A systematic review of empirical evidence. Autism research, 14(4), 604–630.

Caucheteux, C., Gramfort, A., & King, J. R. (2022). Deep language algorithms predict semantic comprehension from brain activity. Scientific reports, 12(1), 16327.

Chaure, F. J., Rey, H. G., & Quian Quiroga, R. (2018). A novel and fully automatic spike-sorting implementation with variable number of features. Journal of Neurophysiology, 120(4), 1859–1871. 10.1152/jn.00339.2018

Chrysaitis, N., & Seriès, P. (2023). 10 years of Bayesian theories of autism: A comprehensive review. Neuroscience and biobehavioral reviews, 145, 105022.

Constantinescu, A. O., O’Reilly, J. X., & Behrens, T. E. (2016). Organizing conceptual knowledge in humans with a gridlike code. Science, 352(6292), 1464–1468.

Courellis, H. S., Minxha, J., Cardenas, A. R., Kimmel, D. L., Reed, C. M., Valiante, T. A., … & Rutishauser, U. (2024). Abstract representations emerge in human hippocampal neurons during inference. Nature, 632(8026), 841–849.

Cruse, D. A. (1986). Lexical semantics. Cambridge: Cambridge University Press.

Cunningham, J. P., & Yu, B. M. (2014). Dimensionality reduction for large-scale neural recordings. Nature neuroscience, 17(11), 1500–1509.

Dale, A. M., Fischl, B., & Sereno, M. I. (1999). Cortical Surface-Based Analysis: I. Segmentation and Surface Reconstruction. NeuroImage, 9(2), 179–194. 10.1006/nimg.1998.0395

Devlin, J., Chang, M.-W., Lee, K., Google, K. T., & Language, A. I. (2018). BERT: Pre-training of Deep Bidirectional Transformers for Language Understanding. https://github.com/tensorflow/tensor2tensor

Dickstein, D. P., Pescosolido, M. F., Reidy, B. L., Galvan, T., Kim, K. L., Seymour, K. E., … & Barrett, R. P. (2013). Developmental meta-analysis of the functional neural correlates of autism spectrum disorders. Journal of the American Academy of Child & Adolescent Psychiatry, 52(3), 279–289.

Dijksterhuis, D. E., Self, M. W., Possel, J. K., Peters, J. C., van Straaten, E. C. W., Idema, S., Baaijen, J. C., van der Salm, S. M. A., Aarnoutse, E. J., van Klink, N. C. E., van Eijsden, P., Hanslmayr, S., Chelvarajah, R., Roux, F., Kolibius, L. D., Sawlani, V., Rollings, D. T., Dehaene, S., & Roelfsema, P. R. (2024). Pronouns reactivate conceptual representations in human hippocampal neurons. Science, 385(6716), 1478–1484.

Duff, M. C., & Brown-Schmidt, S. (2012). The hippocampus and the flexible use and processing of language. Frontiers in human neuroscience, 6, 69.

Ebitz, R. B., & Hayden, B. Y. (2021). The population doctrine in cognitive neuroscience. Neuron, 109(19), 3055–3068.

Ebitz, R. B., & Hayden, B. Y. (2021). The population doctrine in cognitive neuroscience.Neuron, 109(19), 3055–3068.

Ebitz, R. B., Sleezer, B. J., Jedema, H. P., Bradberry, C. W., & Hayden, B. Y. (2019). Tonic exploration governs both flexibility and lapses. PLoS computational biology, 15(11), e1007475.

Eichenbaum, H. (2017). The role of the hippocampus in navigation is memory. Journal of neurophysiology, 117(4), 1785–1796.

Eisenreich, B. R., Hayden, B. Y., & Zimmermann, J. (2019). Macaques are risk-averse in a freely moving foraging task. Scientific reports, 9(1), 15091.

Ethayarajh, K. (2019). How contextual are contextualized word representations? Comparing the geometry of BERT, ELMo, and GPT-2 embeddings. arXiv preprint arXiv:1909.00512.

Feldman, H., & Friston, K. J. (2010). Attention, uncertainty, and free-energy. Frontiers in human neuroscience, 4, 215.

Fellbaum, C. (1998). WordNet: An Electronic Lexical Database. The MIT Press; Bradford Books.

Franch, M., Mickiewicz, E. A., Belanger, J. L., Chericoni, A., Chavez, A. G., Katlowitz, K. A., … & Hayden, B. Y. (2025). A vectorial code for semantics in human hippocampus. bioRxiv, 2025-02.

Friston, K. (2010). The free-energy principle: a unified brain theory?. Nature reviews neuroscience, 11(2), 127–138.

Friston, K. J., Stephan, K. E., Montague, R., & Dolan, R. J. (2014). Computational psychiatry: the brain as a phantastic organ. The Lancet Psychiatry, 1(2), 148–158.

Gao, P., Trautmann, E., Yu, B., Santhanam, G., Ryu, S., Shenoy, K., & Ganguli, S. (2017). A theory of multineuronal dimensionality, dynamics and measurement. BioRxiv, 214262.

Goldstein, A., Zada, Z., Buchnik, E., Schain, M., Price, A., Aubrey, B., … & Hasson, U. (2022). Shared computational principles for language processing in humans and deep language models. Nature neuroscience, 25(3), 369–380.

Goldstein, A., Ham, E., Nastase, S. A., Zada, Z., Grinstein-Dabus, A., Aubrey, B., … & Hasson, U. (2022). Correspondence between the layered structure of deep language models and temporal structure of natural language processing in the human brain. BioRxiv, 2022-07.

Goldstein, A., Wang, H., Niekerken, L., Zada, Z., Aubrey, B., Sheffer, T., Nastase, S. A., Gazula, H., Schain, M., Singh, A., Rao, A., Choe, G., Kim, C., Doyle, W., Friedman, D., Devore, S., Dugan, P., Hassidim, A., Brenner, M., … Hasson, U. (2023). Deep speech-to-text models capture the neural basis of spontaneous speech in everyday conversations. 10.1101/2023.06.26.546557

Goldstein, A., Grinstein-Dabush, A., Schain, M., Wang, H., Hong, Z., Aubrey, B., Nastase, S. A., Zada, Z., Ham, E., Feder, A., Gazula, H., Buchnik, E., Doyle, W., Devore, S., Dugan, P., Reichart, R., Friedman, D., Brenner, M., Hassidim, A., … Hasson, U. (2024). Alignment of brain embeddings and artificial contextual embeddings in natural language points to common geometric patterns. Nature Communications, 15(1), 2768. 10.1038/s41467-024-46631-y

Gómez, C., Lizier, J. T., Schaum, M., Wollstadt, P., Grützner, C., Uhlhaas, P., … & Wibral, M. (2014). Reduced predictable information in brain signals in autism spectrum disorder. Frontiers in neuroinformatics, 8, 9.

Gonzalez-Gadea, M. L., Chennu, S., Bekinschtein, T. A., Rattazzi, A., Beraudi, A., Tripicchio, P., … & Ibanez, A. (2015). Predictive coding in autism spectrum disorder and attention deficit hyperactivity disorder. Journal of Neurophysiology, 114(5), 2625–2636.

Grisoni, L., Moseley, R. L., Motlagh, S., Kandia, D., Sener, N., Pulvermüller, F., … & Mohr, B. (2019). Prediction and mismatch negativity responses reflect impairments in action semantic processing in adults with autism spectrum disorders. Frontiers in Human Neuroscience, 13, 395.

Groppe, D. M., Bickel, S., Dykstra, A. R., Wang, X., Mégevand, P., Mercier, M. R., Lado, F. A., Mehta, A. D., & Honey, C. J. (2017). iELVis: An open source MATLAB toolbox for localizing and visualizing human intracranial electrode data. Journal of Neuroscience Methods, 281, 40–48. 10.1016/j.jneumeth.2017.01.022

Gwilliams, L., Marantz, A., Poeppel, D., King, J.-R., 2024. Hierarchical dynamic coding coordinates speech comprehension in the brain. 10.1101/2024.04.19.590280

Harris, G. J., Chabris, C. F., Clark, J., Urban, T., Aharon, I., Steele, S., … & Tager-Flusberg, H. (2006). Brain activation during semantic processing in autism spectrum disorders via functional magnetic resonance imaging. Brain and cognition, 61(1), 54–68.

Hashimoto, T., Yokota, S., Matsuzaki, Y., & Kawashima, R. (2021). Intrinsic hippocampal functional connectivity underlying rigid memory in children and adolescents with autism spectrum disorder: A case–control study. Autism, 25(7), 1901–1912.

Hayden, B. Y., Heilbronner, S. R., & Yoo, S. B. M. (2026). Rethinking the centrality of brain areas in understanding functional organization. Nature Neuroscience, 29(2), 267–278.

Hayden, B. Y., Smith, D. V., & Platt, M. L. (2010). Cognitive control signals in posterior cingulate cortex. Frontiers in human neuroscience, 4, 223.

He, Y., Yuan, M., Chen, J., & Horrocks, I. (2024). Language models as hierarchy encoders. Advances in Neural Information Processing Systems, 37, 14690–14711.

Hotelling, H. (1936). Relations between two sets of variates. Biometrika, 28(3–4), 321–377. 10.1093/biomet/28.3-4.321

Huth, A. G., de Heer, W. A., Griffiths, T. L., Theunissen, F. E., & Gallant, J. L. (2016). Natural speech reveals the semantic maps that tile human cerebral cortex. Nature, 532(7600), 453–458.

Huys, Q. J., Maia, T. V., & Frank, M. J. (2016). Computational psychiatry as a bridge from neuroscience to clinical applications. Nature neuroscience, 19(3), 404–413.

Izenman, A. J. (1975). Reduced-rank regression for the multivariate linear model. Journal of multivariate analysis, 5(2), 248–264.

Jamali, M., Grannan, B., Cai, J., Khanna, A. R., Muñoz, W., Caprara, I., Paulk, A. C., Cash, S. S., Fedorenko, E., & Williams, Z. M. (2024). Semantic encoding during language comprehension at single-cell resolution. Nature, 631(8021), 610–616.

Jenkinson, M., Bannister, P., Brady, M., & Smith, S. (2002). Improved Optimization for the Robust and Accurate Linear Registration and Motion Correction of Brain Images. NeuroImage, 17(2), 825–841.

Johnston, W. J., Fine, J. M., Yoo, S. B. M., Ebitz, R. B., & Hayden, B. Y. (2024). Semi-orthogonal subspaces for value mediate a binding and generalization trade-off. Nature Neuroscience, 27(11), 2218–2230.

Joshi, A., Scheinost, D., Okuda, H., Belhachemi, D., Murphy, I., Staib, L. H., & Papademetris, X. (2011). Unified Framework for Development, Deployment and Robust Testing of Neuroimaging Algorithms. Neuroinformatics, 9(1), 69–84. 10.1007/s12021-010-9092-8

Karkowski, K., Kehl, M. S., Qin, Y., Darcher, A., Karkowski, P., Borger, V., … & Mormann, F. (2025). Semantic Tuning of Single Neurons in the Human Medial Temporal Lobe. bioRxiv, 2025-10.

Katlowitz, K. A., Belanger, J. L., Ismail, T., Chavez, A. G., Chericoni, A., Franch, M., … & Hayden, B. Y. (2025). Attention is all you need (in the brain): semantic contextualization in human hippocampus. bioRxiv, 2025-06.

Katlowitz, K. A., Cole, E. R., Mickiewicz, E. A., Shah, S., Franch, M. C., Adkinson, J., … & Sheth, S. A. (2025). Plasticity and Language in the Anesthetized Human Hippocampus. Biorxiv, 2025-04.

Khanna, A. R., Muñoz, W., Kim, Y. J., Kfir, Y., Paulk, A. C., Jamali, M., … & Williams, Z. M. (2024). Single-neuronal elements of speech production in humans. Nature, 626(7999), 603–610.

Kolibius, L. D., Roux, F., Parish, G., Ter Wal, M., Van Der Plas, M., Chelvarajah, R., Sawlani, V., Rollings, D. T., Lang, J. D., Gollwitzer, S., Walther, K., Hopfengärtner, R., Kreiselmeyer, G., Hamer, H., Staresina, B. P., Wimber, M., Bowman, H., & Hanslmayr, S. (2023). Hippocampal neurons code individual episodic memories in humans. Nature Human Behaviour, 7(11), 1968–1979. 10.1038/s41562-023-01706-6.

Kriegeskorte, N., & Douglas, P. K. (2018). Cognitive computational neuroscience. Nature neuroscience, 21(9), 1148–1160.

Landauer, T. K. (2001). Single representations of multiple meanings in latent semantic analysis. In D. S. Gorfein (Ed.), On the consequences of meaning selection: Perspectives on resolving lexical ambiguity (pp. 217–232). Washington, D. C.: APA Press.

Lawson, R. P., Rees, G., & Friston, K. J. (2014). An aberrant precision account of autism. Frontiers in human neuroscience, 8, 302.

Lawson, R. P., Mathys, C., & Rees, G. (2017). Adults with autism overestimate the volatility of the sensory environment. Nature neuroscience, 20(9), 1293–1299.

Levy, R. (2008). Expectation-based syntactic comprehension. Cognition, 106(3), 1126–1177.

Li, X., Morgan, P. S., Ashburner, J., Smith, J., & Rorden, C. (2016). The first step for neuroimaging data analysis: DICOM to NIfTI conversion. Journal of Neuroscience Methods, 264, 47–56. 10.1016/j.jneumeth.2016.03.001

Lord, C., Elsabbagh, M., Baird, G., & Veenstra-Vanderweele, J. (2018). Autism spectrum disorder. The lancet, 392(10146), 508–520.

Magnotti, J. F., Wang, Z., & Beauchamp, M. S. (2020). RAVE: Comprehensive open-source software for reproducible analysis and visualization of intracranial EEG data. NeuroImage, 223, 117341. 10.1016/j.neuroimage.2020.117341

Maisson, D. J. N., Cervera, R. L., Voloh, B., Conover, I., Zambre, M., Zimmermann, J., & Hayden, B. Y. (2023). Widespread coding of navigational variables in prefrontal cortex. Current Biology, 33(16), 3478–3488.

Maisson, D. J. N., Cash-Padgett, T. V., Wang, M. Z., Hayden, B. Y., Heilbronner, S. R., & Zimmermann, J. (2021). Choice-relevant information transformation along a ventrodorsal axis in the medial prefrontal cortex. Nature communications, 12(1), 4830.

Manning, C. D., Surdeanu, M., Bauer, J., Finkel, J. R., Bethard, S., & McClosky, D. (2014, June). The Stanford CoreNLP natural language processing toolkit. In Proceedings of 52nd annual meeting of the association for computational linguistics: system demonstrations (pp. 55–60).

Manning, C. D., Clark, K., Hewitt, J., Khandelwal, U., & Levy, O. (2020). Emergent linguistic structure in artificial neural networks trained by self-supervision. Proceedings of the National Academy of Sciences, 117(48), 30046–30054.

Mikolov, T., Chen, K., Corrado, G. S., & Dean, J. (2013). Efficient Estimation of Word Representations in Vector Space. International Conference on Learning Representations. https://api.semanticscholar.org/CorpusID:5959482

Minor, G. N., Hannula, D. E., Gordon, A., Ragland, J. D., Iosif, A. M., & Solomon, M. (2023). Relational memory weakness in autism despite the use of a controlled encoding task.Frontiers in Psychology, 14, 1210259.

Mitchell, P., & Ropar, D. (2004). Visuo-spatial abilities in autism: A review. Infant and Child Development: An International Journal of Research and Practice, 13(3), 185–198.

Montague, P. R., Dolan, R. J., Friston, K. J., & Dayan, P. (2012). Computational psychiatry. Trends in cognitive sciences, 16(1), 72–80.

Noel, J. P., Zhang, L. Q., Stocker, A. A., & Angelaki, D. E. (2021). Individuals with autism spectrum disorder have altered visual encoding capacity. PLoS biology, 19(5), e3001215.

Noel, J. P., & Angelaki, D. E. (2023). A theory of autism bridging across levels of description. Trends in Cognitive Sciences, 27(7), 631–641.

Pellicano, E., & Burr, D. (2012). When the world becomes ‘too real’: a Bayesian explanation of autistic perception. Trends in cognitive sciences, 16(10), 504–510.

Phan, L., Tariq, A., Lam, G., Pang, E. W., & Alain, C. (2021). The neurobiology of semantic processing in autism spectrum disorder: an activation likelihood estimation analysis. Journal of autism and developmental disorders, 51(9), 3266–3279.

Piai, V., Anderson, K. L., Lin, J. J., Dewar, C., Parvizi, J., Dronkers, N. F., & Knight, R. T. (2016). Direct brain recordings reveal hippocampal rhythm underpinnings of language processing. Proceedings of the National Academy of Sciences, 113(40), 11366–11371.

Provenza, N. R., Reddy, S., Allam, A. K., Rajesh, S. V., Diab, N., Reyes, G., … & Sheth, S. A. (2024). Disruption of neural periodicity predicts clinical response after deep brain stimulation for obsessive-compulsive disorder. Nature Medicine, 30(10), 3004–3014.

Qela, B., Damiani, S., De Santis, S., Groppi, F., Pichiecchio, A., Asteggiano, C., … & Fusar-Poli, L. (2025). Predictive coding in neuropsychiatric disorders: A systematic transdiagnostic review. Neuroscience & Biobehavioral Reviews, 169, 106020.

Quiroga, R. Q., Reddy, L., Kreiman, G., Koch, C., & Fried, I. (2005). Invariant visual representation by single neurons in the human brain. Nature, 435(7045), 1102–1107. 10.1038/nature03687

Quian Quiroga, R., Kraskov, A., Koch, C., & Fried, I. (2009). Explicit Encoding of Multimodal Percepts by Single Neurons in the Human Brain. Current Biology, 19(15), 1308–1313. 10.1016/j.cub.2009.06.060

Radford, A., Wu, J., Child, R., Luan, D., Amodei, D., & Sutskever, I. (2019). Language Models are Unsupervised Multitask Learners. https://github.com/codelucas/newspaper

Rao, R. P., & Ballard, D. H. (1999). Predictive coding in the visual cortex: a functional interpretation of some extra-classical receptive-field effects. Nature neuroscience, 2(1), 79–87.

Reinhardt, V. P., Iosif, A. M., Libero, L., Heath, B., Rogers, S. J., Ferrer, E., … & Solomon, M. (2020). Understanding hippocampal development in young children with autism spectrum disorder. Journal of the American Academy of Child & Adolescent Psychiatry, 59(9), 1069–1079.

Sapey-Triomphe, L. A., Pattyn, L., Weilnhammer, V., Sterzer, P., & Wagemans, J. (2023). Neural correlates of hierarchical predictive processes in autistic adults. Nature Communications, 14(1), 3640.

Schapiro, A. C., Turk-Browne, N. B., Botvinick, M. M., & Norman, K. A. (2017). Complementary learning systems within the hippocampus: a neural network modelling approach to reconciling episodic memory with statistical learning. Philosophical Transactions of the Royal Society B: Biological Sciences, 372(1711).

Schneebeli, M., Haker, H., Rüesch, A., Zahnd, N., Marino, S., Paolini, G., … & Stephan, K. E. (2022). Disentangling “Bayesian brain” theories of autism spectrum disorder. Medrxiv, 2022-02.

Schneider, F., & Blank, H. (2026). Sensory sharpening and semantic prediction errors unify competing models of predictive processing in human speech comprehension. PLoS Biology, 24(1), e3003588.

Schrimpf, M., Blank, I. A., Tuckute, G., Kauf, C., Hosseini, E. A., Kanwisher, N., … & Fedorenko, E. (2021). The neural architecture of language: Integrative modeling converges on predictive processing. Proceedings of the National Academy of Sciences, 118(45), e2105646118.

Semedo, J. D., Zandvakili, A., Machens, C. K., Yu, B. M., & Kohn, A. (2019). Cortical areas interact through a communication subspace. Neuron, 102(1), 249–259.

Shohamy, D., & Turk-Browne, N. B. (2013). Mechanisms for widespread hippocampal involvement in cognition. Journal of Experimental Psychology: General, 142(4), 1159.

Tang, J., LeBel, A., Jain, S., & Huth, A. G. (2023). Semantic reconstruction of continuous language from non-invasive brain recordings. Nature Neuroscience, 26(5), 858–866. 10.1038/s41593-023-01304-9

Van Boxtel, J. J., & Lu, H. (2013). A predictive coding perspective on autism spectrum disorders. Frontiers in psychology, 4, 19.

Van de Cruys, S., Evers, K., Van der Hallen, R., Van Eylen, L., Boets, B., De-Wit, L., & Wagemans, J. (2014). Precise minds in uncertain worlds: predictive coding in autism. Psychological review, 121(4), 649.

van Laarhoven, T., Stekelenburg, J. J., Eussen, M. L., & Vroomen, J. (2020). Atypical visual-auditory predictive coding in autism spectrum disorder: Electrophysiological evidence from stimulus omissions. Autism, 24(7), 1849–1859.

Vaswani, A., Brain, G., Shazeer, N., Parmar, N., Uszkoreit, J., Jones, L., Gomez, A. N., Kaiser, Ł., & Polosukhin, I. (2017). Attention Is All You Need.

Vig, J., 2019. A Multiscale Visualization of Attention in the Transformer Model, in: Costa-jussà, M.R., Alfonseca, E. (Eds.), Proceedings of the 57th Annual Meeting of the Association for Computational Linguistics: System Demonstrations. Association for Computational Linguistics, Florence, Italy, pp. 37–42. 10.18653/v1/P19-3007

Viganò, S., & Piazza, M. (2021). The hippocampal-entorhinal system represents nested hierarchical relations between words during concept learning. Hippocampus, 31(6), 557–568.

Vogindroukas, I., Stankova, M., Chelas, E. N., & Proedrou, A. (2022). Language and speech characteristics in autism. Neuropsychiatric disease and treatment, 2367–2377.

Voloh, B., Maisson, D. J. N., Cervera, R. L., Conover, I., Zambre, M., Hayden, B., & Zimmermann, J. (2023). Hierarchical action encoding in prefrontal cortex of freely moving macaques. Cell Rep. 42, 113091.

Wachtel, L. E., Escher, J., Halladay, A., Lutz, A., Satriale, G. M., Westover, A., & Lopez-Arvizu, C. (2024). Profound autism: An imperative diagnosis. Pediatric Clinics, 71(2), 301–313.

Wang, X. J., & Krystal, J. H. (2014). Computational psychiatry. Neuron, 84(3), 638–654.

Wang, M. Z., & Hayden, B. Y. (2021). Latent learning, cognitive maps, and curiosity. Current Opinion in Behavioral Sciences, 38, 1–7.

Weissbart, H., Kandylaki, K. D., & Reichenbach, T. (2020). Cortical tracking of surprisal during continuous speech comprehension. Journal of cognitive neuroscience, 32(1), 155–166.

Weissbart, H., Martin, A.E., 2024. The structure and statistics of language jointly shape cross-frequency neural dynamics during spoken language comprehension. Nat. Commun. 15, 8850. 10.1038/s41467-024-53128-1

Whittington, J. C., Muller, T. H., Mark, S., Chen, G., Barry, C., Burgess, N., & Behrens, T. E. (2020). The Tolman-Eichenbaum machine: unifying space and relational memory through generalization in the hippocampal formation. Cell, 183(5), 1249–1263.

Willems, R. M., Frank, S. L., Nijhof, A. D., Hagoort, P., & Van den Bosch, A. (2016). Prediction during natural language comprehension. Cerebral cortex, 26(6), 2506–2516.

Wu, B., & Pillow, J. (2025). Reduced rank regression for neural communication: a tutorial for Neuroscientists.

Yacoub, E, Grier, M. D., Auerbach, E. J., Lagore, R. L., Harel, N., Adriany, G., Zilverstand A., Hayden, B., Y., Heilbronner, S. R., Ugurbil, K., and Zimmermann, J. (2020). Ultra-high field (10.5 T) resting state fMRI in the macaque. Neuroimage 223: 117349.

Yan, X., Krishna, A., Arsdel, K. V., Gautam, I., Kim, B., Shrivastava, A., … & Sheth, S. A. (2025). Shared neural geometries for bilingual semantic representations. bioRxiv, 2025-11.

Yang, A. I., Wang, X., Doyle, W. K., Halgren, E., Carlson, C., Belcher, T. L., Cash, S. S., Devinsky, O., & Thesen, T. (2012). Localization of dense intracranial electrode arrays using magnetic resonance imaging. NeuroImage, 63(1), 157–165. 10.1016/j.neuroimage.2012.06.039

Zada, Z., Goldstein, A., Michelmann, S., Simony, E., Price, A., Hasenfratz, L., Barham, E., Zadbood, A., Doyle, W., Friedman, D., Dugan, P., Melloni, L., Devore, S., Flinker, A., Devinsky, O., Nastase, S. A., & Hasson, U. (2024). A shared model-based linguistic space for transmitting our thoughts from brain to brain in natural conversations. Neuron, 112(18), 3211–3222.e5. 10.1016/j.neuron.2024.06.025

Zhu, H., Franch, M., Mickiewicz, E., Belanger, J., Cowan, R. L., Katlowitz, K., … & Hayden, B. Y. (2026). Semantic axes in the brain support analogical representations. bioRxiv, 2026-01.

